# A TCF4/BRD4-dependent regulatory network confers cross-resistance to targeted and immune checkpoint therapy in melanoma

**DOI:** 10.1101/2022.08.11.502598

**Authors:** Joanna Pozniak, Dennis Pedri, Ewout Landeloos, Yannick Van Herck, Asier Antoranz, Panagiotis Karras, Ada Nowosad, Samira Makhzami, Greet Bervoets, Michael Dewaele, Lukas Vanwynsberghe, Sonia Cinque, Sam Kint, Katy Vandereyken, Thierry Voet, Frank Vernaillen, Wim Annaert, Diether Lambrechts, Veerle Boecxstaens, Joost van den Oord, Francesca Bosisio, Eleonora Leucci, Florian Rambow, Oliver Bechter, Jean-Christophe Marine

**Author notes:** These authors have contributed equally to this work (Joanna Pozniak, Dennis Pedri, Ewout Landeloos, Yannick Van Herck).

## Abstract

Primary resistance drastically limits the clinical success of immune checkpoint blockade (ICB) in melanoma. Resistance to ICB may also develop when tumours relapse after targeted therapy. To identify cancer cell-intrinsic mechanisms driving resistance to ICB, we generated single-cell RNA-sequencing (scRNA-seq) data from a prospective longitudinal cohort of patients on ICB therapy, including an early time point obtained after only one cycle of treatment. Comparing these data with murine scRNA-seq datasets, we established a comprehensive view of the cellular architecture of the treatment-naïve melanoma ecosystem, and defined 6 evolutionarily conserved melanoma transcriptional metaprograms (Melanocytic or MEL, Mesenchymal-like or MES, Neural Crest-like, Antigen Presentation, Stress (hypoxia response) and Stress (p53 response)). Spatial multi-omics revealed a non-random geographic distribution of cell states that is, at least partly, driven by the tumour microenvironment. The single-cell data allowed unambiguous discrimination between melanoma MES cells and cancer-associated fibroblasts both *in silico* and *in situ*, a long-standing challenge in the field. Importantly, two of the melanoma transcriptional metaprograms were associated with divergent clinical responses to ICB. While the Antigen Presentation cell population was more abundant in tumours from patients who exhibited a clinical response to ICB, MES cells were significantly enriched in early on-treatment biopsies from non-responders, and their presence significantly predicted lack of response. Critically, we identified TCF4 (E2-2) as a master regulator of the MES program and suppressor of both MEL and Antigen Presentation programs. Targeting *TCF4* expression in MES cells either genetically or pharmacologically using a bromodomain inhibitor increased immunogenicity and sensitivity to targeted therapy. This study describes an increasingly complex melanoma transcriptional landscape and its rapid evolution under ICB. It also identifies a putative biomarker of early response to ICB and an epigenetic therapeutic strategy that increases both immunogenicity of ICB-refractory melanoma and their sensitivity to targeted therapy.

## Introduction

Despite several breakthroughs in the field, metastatic melanoma (MM) continues to be a major clinical challenge^1, 2^. Although treatment outcomes have substantially improved since the introduction of immune checkpoint blockade (ICB)^3, 4^ approximately half of the MM patients do not gain any durable survival benefit. One of the key challenges is therefore to elucidate why ICB therapies, such as anti-PD-1, anti-CTLA-4 or their combination, are effective in some, but not all, patients, and ultimately identify rational therapeutic (combination) strategies that overcome resistance.

Tumour-extrinsic and intrinsic mechanisms can drive resistance to ICB^5, 6^. For instance, tumour mutational burden has been associated with ICB response through increased neoantigen formation, bolstering immunogenicity^7, 8^. Inactivating mutations in genes encoding components of the antigen processing and/or presentation machinery (e.g., MHC class I, B2-microglobulin) can lead to ICB resistance. Similarly, tumours with inactivating mutations in JAK1/JAK2 are associated with loss of interferon responsiveness, and thereby resistance to PD-1 blockade^9,10^.

In addition, there is increasing evidence that melanoma cells can adopt a variety of phenotypic states through nongenetic reprogramming, and thereby exhibit different sensitivities to cancer treatments, including ICB^11^. Dedifferentiation of melanoma cells was previously described as such a nongenetic mechanism that drives immune escape and resistance to adoptive T cell transfer^12, 13^. Based on bulk RNA-seq data analyses of anti-PD-1 treated melanoma patients, with samples collected at baseline and upon progression, it was further proposed that dedifferentiation may also be a mechanism driving resistance to ICB^14^. Deconvolution of additional bulk RNA-seq datasets and immunostaining further confirmed the enrichment of a dedifferentiated (NGFR^high^) Neural-Crest-like program in tumours associated with immune-exclusion^15^ and resistance to immunotherapy^16^. Mechanistically, dedifferentiation was proposed to dampen response to ICB due to a decrease in expression and/or presentation of melanocytic antigens^12, 13^, MHC class I downregulation^17^, and secretion of the neurotrophic factor BDNF, which contributes to resistance to antigen-specific T cells^15^. Consistent with these findings, the innate PD-1-inhibitor resistance (IPRES) signature (which was defined based on the analysis of bulk RNA-seq data and includes 26 gene signatures associated with dedifferentiation) was associated with poor response to anti-PD-1 in pre-treatment biopsies^14^. However, such an association could not be established in other melanoma cohorts^18, 19^. The difficulty in identifying reliable predictive information at baseline using bulk transcriptomic data was further exemplified by the lack of reproducibility when predicting response to ICB using IMPRES^16, 20^, yet another gene expression signature derived from bulk RNA-seq datasets^21^. Bulk longitudinal analyses later confirmed that a robust pre-treatment biomarker is unlikely to capture the heterogenous nature of cancer and/or anticipate the rapid evolution of tumour phenotypes under ICB therapy^17^.

It has therefore become evident that understanding resistance to ICB requires single-cell resolution and temporal dissection of the entire cellular architecture of the melanoma ecosystem. Using scRNA-seq, a MYC-driven malignant gene expression signature associated with immune evasion and T-cell exclusion was recently identified^22^. Although very informative, this study was limited by the recovery of a relatively small number of malignant cells and absence of patient-matched samples across both time points. In addition, only one responder was identified in the discovery cohort. It is important to note that, equal to the previous study by the same group^23^, most biopsies in this study originated from patients with prior exposure to diverse treatments. Therefore, a comprehensive view of the cellular architecture of the treatment-naïve melanoma ecosystem, and in particular of its transcriptomic landscape, and its evolution under ICB therapy is still lacking.

## Results

### Portraying the treatment-naive human melanoma transcriptomic landscape

To dissect the cellular composition of the human melanoma ecosystem and study how it evolves under ICB, we set up a unique prospective longitudinal study including treatment naïve stage III/IV (AJCC 8^th^ edition) melanoma patients receiving anti-PD-1 based therapy (anti-PD-1 monotherapy (nivolumab): n=17; anti-PD-1 and anti-CTLA4 combination therapy (ipilimumab + nivolumab): (n=6) SPECIAL trial; UZ/KU Leuven #S62275). Cutaneous, subcutaneous or lymph node metastases were biopsied before initiation of therapy (before treatment; BT). Subsequently, a second tumour biopsy was collected right before the administration of the second ICB treatment cycle (early on-treatment; OT). We obtained patient- and lesion-matched biopsies across both time points for 20 patients. Part of the obtained material was preserved for routine pathological assessment, multiplex immunohistochemistry (mIHC), multiplex RNA fluorescence *in situ* hybridization (mFISH) and untargeted spatial transcriptomics. The remaining tissue was processed for single-cell transcriptome profiling (Figure 1A). Demographic, clinical, histopathological and genetic information was collected at baseline. Patients with unresectable disease were stratified as responders (complete remission, partial remission) and non-responders (stable disease, progressive disease) based on RECISTv1.1. best overall response, whereas patients treated with curative intent were stratified according to pathological response assessment at tumour resection^24^.

**Figure 1:**
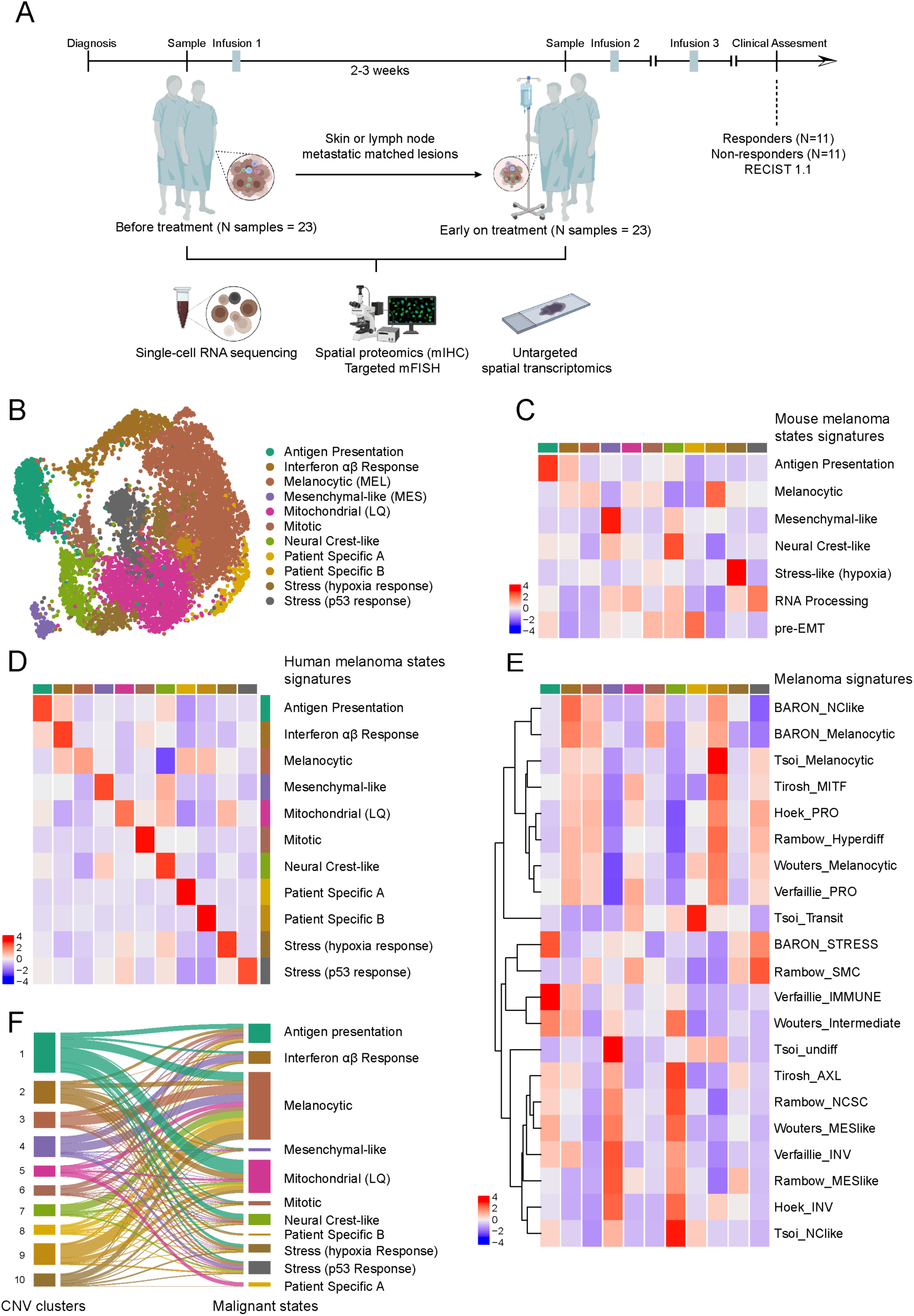
Refining the human melanoma transcriptomic landscape. **A,** Study design, including timing of sample collection and processing methodologies. **B,** Uniform Manifold Approximation and Projection (UMAP) of malignant cells, containing 11 functionally annotated clusters. **C,** AUCell scores of the functionally enriched marker genes per malignant mouse state (acquired from Karras *et. al*^32^), averaged and plotted across the malignant human states. **D,** AUCell scores of the top 100 marker genes of the malignant states averaged and replotted across each state. **E,** AUCell scores of various previously published melanoma signatures averaged and plotted across each state. **F,** Alluvial plot intersecting each cell, represented in clusters based on CNV patterns and in the transcriptomic states.

In total, > 59K single cells passed our quality control requirements (see methods). To dissect the cellular composition of the melanoma tumour microenvironment (TME), we first measured the activity of previously described stromal, immune and malignant/melanoma gene sets^22^ and assigned each cell from the unsupervised clusters to one of these three compartments (Supplemental Figure S1A-C). Importantly, each of these compartments could be detected in all lesions, irrespective of their metastatic site of origin (Supplemental Figure S1D).

We further refined our malignant cell annotation pipeline to filter out cells that do not exhibit large-scale genomic rearrangements (Supplemental Figure S1E), and do not harbour a high score for the malignant signature described by Jerby-Arnon and colleagues^22^ or for the “Melanoma Signature”, which we generated to discriminate between dedifferentiated melanoma cells and cancer-associated fibroblasts (Supplemental Figure S1E,F and Supplemental Table S1; see methods and below). We also excluded cells expressing the immune cell marker gene *PTPRC* (CD45).

As previously reported for melanoma^22, 23, 25^ and other human cancers^26, 27^, dimension reduction visualisation showed a clear separation of malignant cells per patient, thereby impeding identification of shared transcriptomic states (Supplemental Figure S2A). This was partly overcome by regressing for patient ID and integrating the data using Harmony^28^ (Supplemental Figure S2B,C). Silhouette^29^ scores were measured to identify the optimal clustering resolution (Supplemental Figure S2D), which we initially set to 12 Seurat malignant clusters (Supplemental Figure S2E). Differential gene expression (DEG) analysis resulted in characteristic gene lists for each cluster (Supplemental Table S2). This analysis prompted us to merge cluster 0 and cluster 2 as they exhibited a similar enrichment for ribosomal genes, thus yielding 11 distinct malignant clusters.

For an in-depth characterisation of the treatment-naïve melanoma ecosystem, we performed another DEG analysis focusing on the untreated samples and interpreted the differentially expressed gene lists (Supplemental Figure S3A and Supplemental Table S3) using EnrichR^30^ across multiple databases (Supplemental Table S4). Gene regulatory modules were also defined for each cluster using SCENIC^31^ (Supplemental Figure S3B and Supplemental Table S5).

For the functional annotation of malignant clusters (Figure 1B) we relied both on the interpretation of DEG lists, with various gene set enrichment tools (Supplemental Table S4), and prior biological knowledge acquired through analysis of a scRNA-seq dataset from Tyr::NRasQ61K/°;Ink4a-/-mouse tumours^32^. Unsupervised clustering of these mouse lesions identified 7 distinct melanoma cell states, which we named Melanocytic, Mesenchymal-like, Neural Crest-like, Stress (hypoxia), RNA-processing, Stem-like (pre-EMT) and Antigen Presenting cell states. 6 of these 7 murine melanoma states overlapped with cellular states identified in the human lesions (Figure 1C). These included the Melanocytic (MEL), Mesenchymal-like (MES), Antigen Presentation, Neural Crest-like, Stress (hypoxia response) cell states. The murine RNA processing state largely overlapped with the human Stress (p53 response) state. Among the previously described marker genes *NGFR* was identified in the Neural-Crest-like state, *VEGFA* was highly expressed in Stress (hypoxia response) cells, and collagen genes (i.e. *COL5A1*) in MES cells (Supplemental Figure S3A). HLA class I and II and other genes involved in antigen processing and presentation, such as *TAP1*, *B2M* and *NLRC5,* were identified as discriminative markers of the Antigen Presentation cell population (Supplemental Figure S3A and Supplemental Table S3). Antigen Presentation cells also expressed many of the canonical interferon-stimulated genes, including among others STAT1-regulated genes (Supplemental Figure S3B and Supplemental Table S3).

The cross-species comparison also highlighted five human-specific cell states: an Interferon Alpha/Beta Response state expressing interferon type I responsive genes (i.e. *IFI6*, *IFI27*, *IRF7*), but not genes involved in antigen processing and presentation. A Mitotic state, which expressed high levels of *MKI67* and *TOP2A*, was also identified, as well as a Mitochondrial state, which exhibited high mitochondrial gene expression and showed no consistent pathway enrichment. We annotated this latter cell state as a “low quality” (LQ) malignant cell cluster. Mitotic and Mitochondrial (LQ) cell clusters are both routinely identified in human tumour biopsy samples^33, 34^. Finally, two patient-specific clusters (Patient-specific A and Patient-specific B), which did not exhibit any specific recognizable functional features, emerged at this level of resolution. Since these clusters were only detected in individual patients, we postulated that they may be driven by specific genetic alterations. Note that while the murine pre-EMT stem-like state did not emerge as an independent cluster, supervised analysis highlighted human melanoma cells from different patients residing in this state^32^.

Using the gene signature of each state we calculated signature scores and visualized each score per state (Figure 1D). While the cellular heterogeneity of melanoma cells broadly aligned with the 11 cell states, a substantial fraction of cells was not exclusively constrained to these states, indicating that melanoma cells can manifest multiple and/or overlapping phenotypes.

The MITF rheostat model predicts that melanoma cell state identity is regulated by the activity of the MITF transcription factor (TF)^11^. The proliferative/melanocytic and dedifferentiated invasive/mesenchymal-like states exhibit high and low MITF activity, respectively. Measuring the activity of these gene expression programs^35, 36^ across all malignant cells confirmed that cells with varying MITF activity co-exist in drug-naïve human metastatic melanoma lesions (data not shown). As expected, the Neural Crest-like and MES states were the most dedifferentiated states. We also measured the activity of a series of previously published melanoma transcriptional cell states^37–40^ identified in various cellular and/or *in vivo* model systems to confirm their identity and/or presence in clinical samples^22^ (Figure 1E and Supplemental Table S6).

Next, we grouped all malignant cells in distinct Copy Number Variation (CNV) genomic clusters using the inferred CNV profiles described in Supplemental Figure S1E. An alluvial plot was used to connect the genomic and transcriptomic clusters for each cell (Figure 1F). Except for cells from the patient-specific clusters A and B, all transcriptional clusters were fed with cells from different genomic clusters.

Moreover, we were able to retrieve all the metaprograms in most of the samples and we did not observe any association between the abundance of a particular melanoma cell state and a specific oncogenic driver mutation (Supplemental Figure S4C, D).

Together these analyses identified several recurrent and evolutionarily conserved transcriptional metaprograms in melanoma, which do not appear to be driven by genetic intra-tumour heterogeneity, but instead are likely to be specified by cues emanating from the tumour microenvironment.

### Spatially mapping of melanoma cell state diversity

To gain insights into the spatial organization of the various melanoma cell states in drug-naïve lesions, we performed untargeted spatially resolved transcriptomics on selected samples (n=6; S1 to S6) from our patient cohort, using the 10X Genomics Visium platform. Each section was annotated by a pathologist based on the morphology of the associated haematoxylin and eosin (H&E) staining. Regions were labelled as either malignant, stromal or immune.

On the Visium platform, multiple (often different) cell types contribute to the transcription profile of each capture area or spot (up to 20 cells/spot). Therefore, to properly capture the nuances of the molecular profile of each patient, and to not risk quenching weak signals, each slide/patient was analysed separately^41^. To spatially resolve the malignant cell states, the spatial transcriptomics data was integrated with the scRNA-seq data using Seurat-v3 anchor-based (CCA) integration^42^ and CellTrek^43^ deconvolution methods (Figure 2A and Supplemental Figure S5A). The Seurat anchor-based integration confirmed that spots in cancer regions were highly enriched for the malignant signature and spots falling outside of the malignant areas were enriched with stromal and/or immune cells (data not shown). These findings were considered as affirmative of our mapping’s validity.

**Figure 2:**
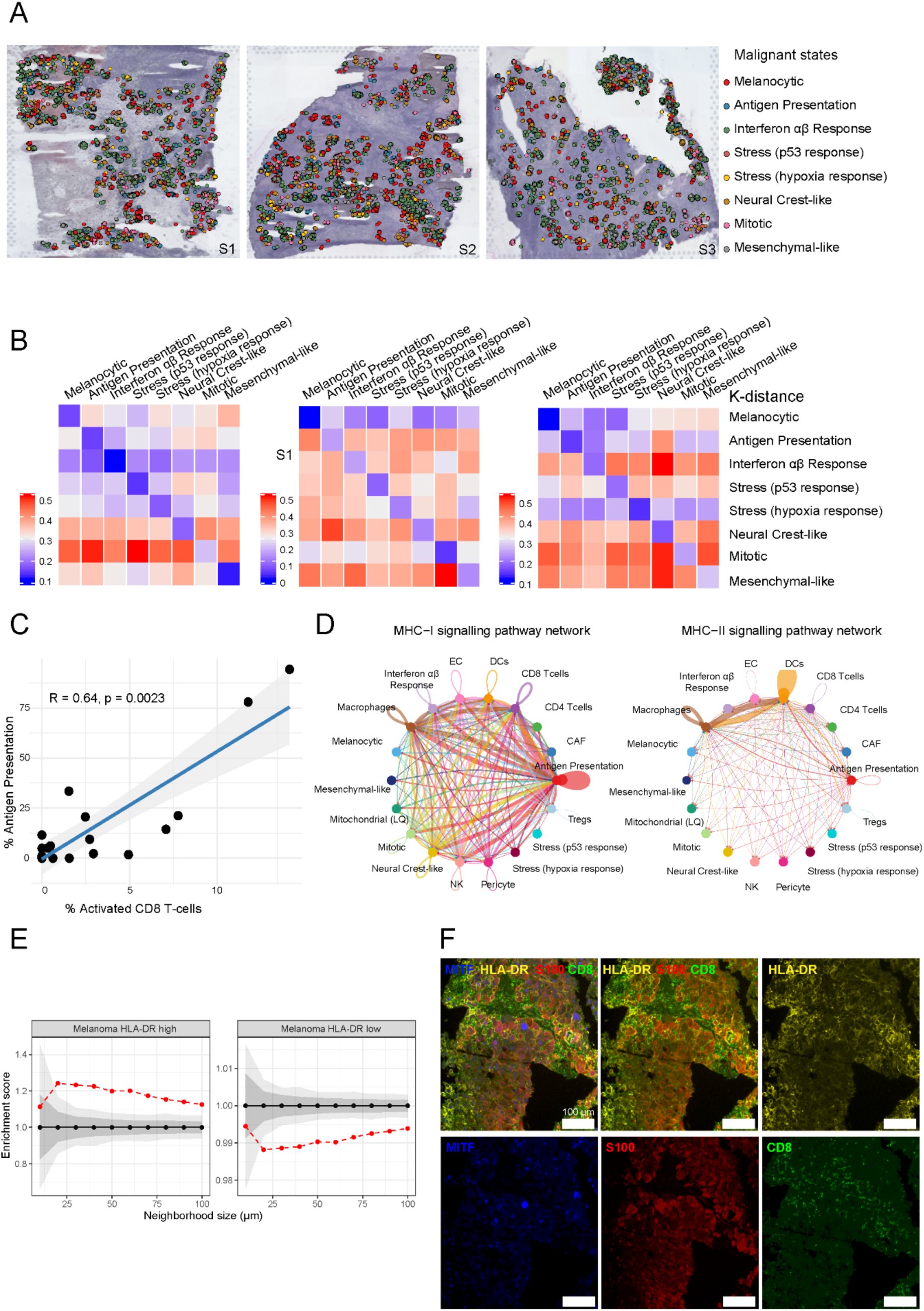
Treatment-naïve melanoma ecosystem mapped spatially. **A,** Spatial transcriptomics on three representative treatment naïve metastatic samples. Shown are malignant spots annotated per state. **B,** Heatmaps of the k-distance calculated between the query state and every other cell state (rows). Note that as the k-distance metric is not normalized to the number of cells, comparisons can only be made within rows. **C,** Correlation of the percentage of the Antigen Presentation state within the tumour compartment with the percentage of activated CD8+ T cells within the immune compartment for each scRNA-seq sample. **D,** Cell-cell interaction prediction from the scRNA-seq data between cells from the malignant and tumour immune microenvironment compartments, via MHC class I and II molecules. The width of the line linking cells indicates the probability strength. **E,** Neighbourhood analysis of the HLA-DR high and low expressing melanoma cells with CD8 T cells in treatment naïve samples (n = 10) from the MILAN data. X-axis: interaction distance considered for neighbourhood analysis (μm). Y-axis: interaction score (positive values indicate interaction, negative avoidance). **F,** Representative image of a clinical biopsy in which HLA-DR high, but not HLA-DR low, melanoma cells co-localise with the CD8+ immune infiltrate.

These spatial transcriptomics data further confirmed the existence of the above-described melanoma cell states, including the two stress states (hypoxia response and p53 response), among cancer cells in the absence of dissociation for scRNA-seq. Zooming further into the cancer regions revealed that melanoma transcriptional metaprograms are not randomly distributed but tend to co-occur in spatially restricted clusters (Figure 2A-B and Supplemental Figure S5A-B). The differential spatial distribution of the malignant cell states can be further highlighted through co-localisation analyses. We found that, as opposed to Mesenchymal-like cells, cells harbouring the Neural Crest-like state preferentially co-occurred with Antigen presentation cells (Supplemental Figure S5C).

Using the scRNA-seq data, we further established a significant positive correlation between the percentage of cells harbouring the Antigen Presentation state and activated CD8^+^ T cells (Figure 2C). Quantitative inference and analysis of intercellular communication networks predicted a functional interaction between these two cell types through engagement of the MHC class I and II signalling pathways (Figure 2D). Consistently, the Antigen Presentation cell state was enriched in lesions with an immune inflamed, often referred to as brisk, phenotype (Supplemental Figure S5D). To further establish a spatial relationship between these two cell types, we performed mIHC using multiple iterative labelling by antibody neodeposition^44^ (MILAN) on treatment naïve melanoma samples (n=10). Neighbourhood analysis confirmed enrichment of melanoma cells positive for the MHC class II marker HLA-DR in the proximity of CD8^+^ T cells (Tcy; Figure 2E,F). In contrast, HLA-DR-negative melanoma cells and CD8^+^ T cells occurred in mutually exclusive regions.

Together, these findings indicate that the transcriptomic heterogeneity of melanoma is spatially organized within the tumour architecture and is, at least partly, driven by heterotypic cellular interactions with the tumour microenvironment. For instance, by integrating signalling predictions with cellular proximity, the data suggest that the melanoma Antigen Presentation cell population emerge by direct interaction with immune cells (i.e. CD8^+^ T cells).

### Unambiguous detection of melanoma MES cells

Similar to epithelial cancer cells that have undergone Epithelial-to-Mesenchymal Transition (EMT), melanoma cells that acquired a mesenchymal-like/dedifferentiated phenotype closely resemble normal mesenchymal cells and cancer associated fibroblasts (CAFs) in particular^11, 45, 46^. Findings concerning EMT through the analysis of bulk-level expression data from human tumours have therefore been confounded by the presence of CAFs^47^. Moreover, identification of coherent and specific marker gene sets that distinguish CAFs and malignant cells that underwent EMT has been a major challenge in the field. Recently, an approach for decoupling the mesenchymal expression profiles of cancer cells and CAFs leveraging scRNA-seq datasets was developed and applied to various epithelial cancers^47^. Unexpectedly, there was no clear evidence for a full EMT malignant state, indicating that this state either does not exist, or is extremely rare and/or transient. Instead, cancer cell-specific partial EMT (pEMT) programs that are distinct from CAF signatures were defined. Even more surprisingly, pEMT was not associated with any specific clinical features across cancers, thereby indicating that the clinical relevance of pEMT expression programs may be highly context-specific. Our single cell analyses did, however, identify melanoma cells expressing a full MES program in both human and mouse^32^ datasets. In our human dataset, the 50 most abundantly expressed genes in MES cells were remarkably almost all highly expressed in CAFs (Figure 3A, left panel and Supplemental Figure S6A, B). In order to define a melanoma-specific Mesenchymal-like gene expression signature, we established a list of the 50 most differentially expressed genes between melanoma MES cells and CAFs (Figure 3A right panel and Supplemental Table S7). Several of these genes including *CDH19* and *S100A1*, which we termed Minimal Lineage Genes (MLGs), were identified in both mouse and human MES signatures and were indeed highly and selectively expressed in MES cells (Figure 3B). Importantly, expression of these genes was higher than *MITF* and *SOX10*, two melanoma markers known to be expressed at very low to undetectable levels in the dedifferentiated MES cells. In contrast, whereas stromal genes like *THY1*, *LUM* and *DCN* were expressed at higher levels in CAFs than in MES cells, several markers including the basic helix-loop-helix TF *TCF4* (also known as ITF2 or E2-2) were instead expressed at comparable levels in both cell types (Figure 3C). Note that, consistent with previous findings^48^, *TCF4* expression was also detected in endothelial (ECs) and plasmacytoid dendritic cells (pDCs; data not shown). Measuring expression of these genes in all melanoma states revealed that while *CDH19* and *S100A1* were expressed in all of them (including MES cells), *TCF4* and other stromal genes were selectively expressed in melanoma MES cells (Figure 3D). We conclude that the MLGs provide the field with a unique tool to unambiguously discriminate between dedifferentiated/mesenchymal-like melanoma cells and CAFs in both mouse and human single-cell datasets.

**Figure 3:**
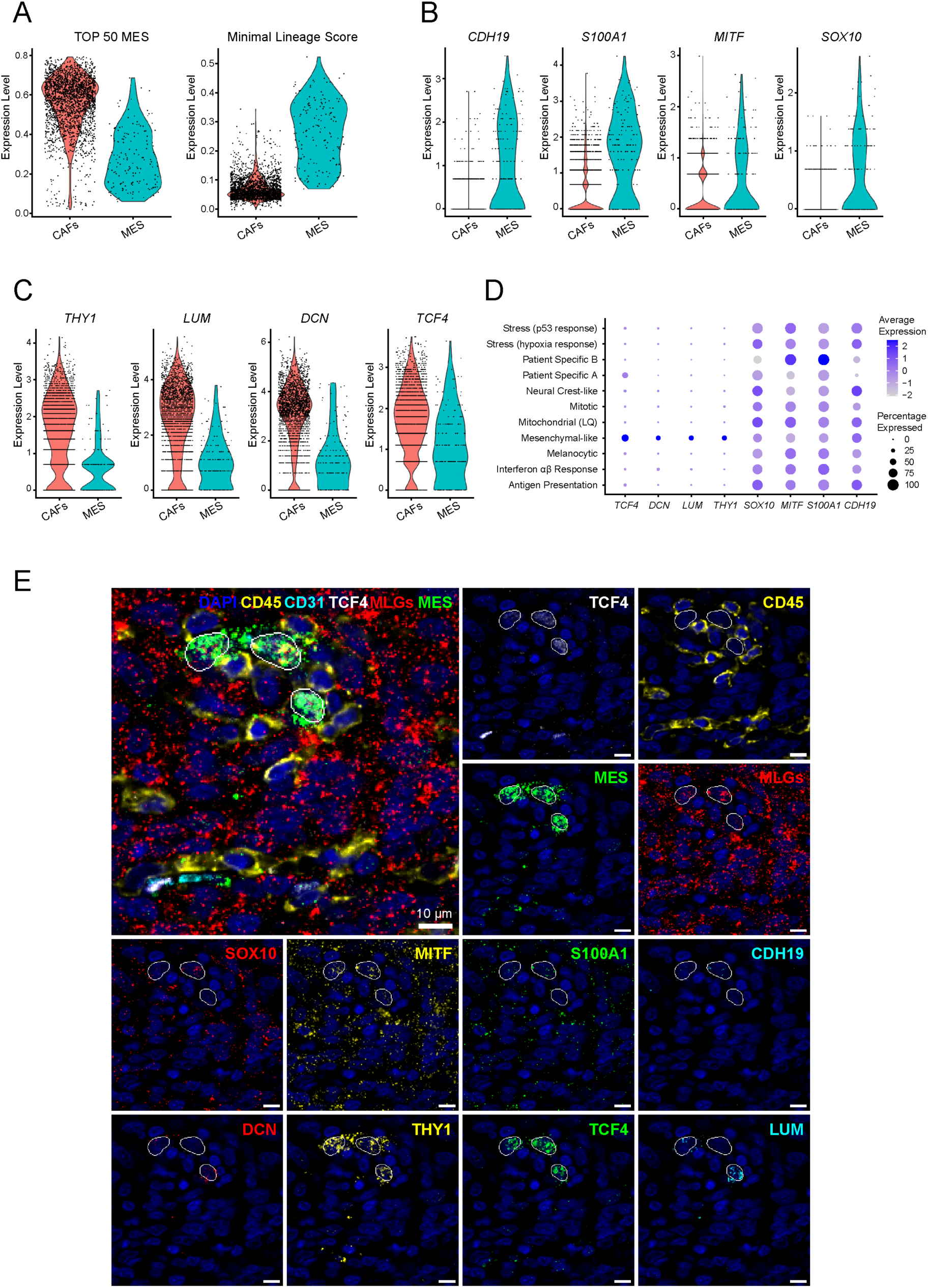
Identification and *in situ* mapping of melanoma MES cells. **A,** AUCell score of the top 50 marker genes of the MES state identified by testing MES vs all other malignant states (left), and Minimal Lineage Genes signature (MLGs) identified by testing MES versus CAFs (right), plotted for CAFs and MES cells. **B,** Expression of four selected MLG marker genes in CAFs and MES cells. **C,** Expression of four selected MES marker genes in CAFs and MES cells. **D,** Expression of the marker genes from B and C across all malignant cell states. **E,** Combined mIHC and mFISH image of a representative treatment naïve lymph node metastasis. CD45, CD31 and TCF4 (white) protein stains are shown, whereas FISH of the four selected genes for the MLGs (*MITF*, *SOX10*, S100A1 and *CDH19; red)* and MES state (*DCN*, *TCF4*, *THY1* and *LUM*; green) are combined (top). A colour-split of each four genes is shown (bottom). Nuclei of Mesenchymal-like cells co-expressing MLGs and MES genes are segmented based on DAPI staining (dark blue).

We next sought to devise a method for the detection and mapping of Mesenchymal-like cells *in situ*. Because expression of the MLGs is relatively low in MES cells, we opted to combine a highly sensitive detection mFISH method (RNAscope) with a multiplexed protein staining assay CODEX for CO-Detection by indEXing^49^. Guided by our scRNA-seq data, we designed a mFISH panel selecting the most discriminatory MLGs (*S100A1* and *CDH19*) and MES (*THY1*, *DCN*, *LUM*) markers to complement a broad panel of melanoma, immune and stromal protein markers (see methods for a full list). We included the pan-mesenchymal marker TCF4, as well as MITF and SOX10, in both our protein and RNA panels. We first tested the method on a selected treatment naïve melanoma lesion. MES cells were identified by co-staining of MLGs (CDH19 and S100A1) and melanoma markers (MITF and SOX10) with MES markers (DCN, THY1, LUM and TCF4) within the CD45-negative cell population. Instead, CAFs were positive for the MES markers and negative for MLGs. Other melanoma subpopulations were positive for the MEL and MLG markers and negative for the MES markers (Figure 3E). Note that pDCs were identifiable as CD45+ CD31-MES-MLGs-, and ECs were CD45-CD31+ MES+ MLGs-(data not shown). To further validate our method, we selected another melanoma sample that was particularly rich in melanoma MES cells and harboured the BRAF^V600E^ mutation. We stained adjacent sections with our combined multiplex IHC/FISH protocol, and with an antibody directed against BRAF^V600E^ mutation. As expected, this sample contained a very high proportion of cells identified as melanoma MES cells (Supplemental Figure S6C). These same cells also stained for the BRAFV^600E^-specific antibody, thus further confirming their malignant origin.

Together, these data provide methodologies to unambiguously identify true melanoma MES cells in both scRNA-seq datasets and on tissue sections, and firmly establish the presence of these cells in human treatment-naïve melanoma lesions.

### MES cells are enriched in early on-treatment melanomas refractory to ICB

Having established the cellular architecture of the drug-naïve melanoma ecosystem and the necessary tools for the unambiguous annotation of all malignant cell states, we next studied how one cycle of ICB therapy may remodel the melanoma transcriptional landscape. There were no overall differences in the proportion of the various melanoma cell states between the BT and OT time points grouping responding and non-responding patients. (Supplemental Figure S7A). Interestingly, however, two of the melanoma cell states, namely the MES and Antigen Presentation states were associated with divergent clinical responses (Figure 4A). Whereas the Antigen Presentation cell state was enriched in OT samples from responders (R), the MES cells were significantly enriched in OT samples from non-responders (NR). The trend of increased abundance of Antigen Presentation cells in lesions from R compared to NR was already observed BT, but this difference was further enhanced at the OT time point (Figure 4A). This observation is consistent with previous findings showing that a subset of melanomas, which harbour tumour cells expressing MHC class II (HLA-DR) molecules, are characterized by an increased CD8+ tumour infiltrate and favourable response to anti-PD-1 therapy^50^.

**Figure 4:**
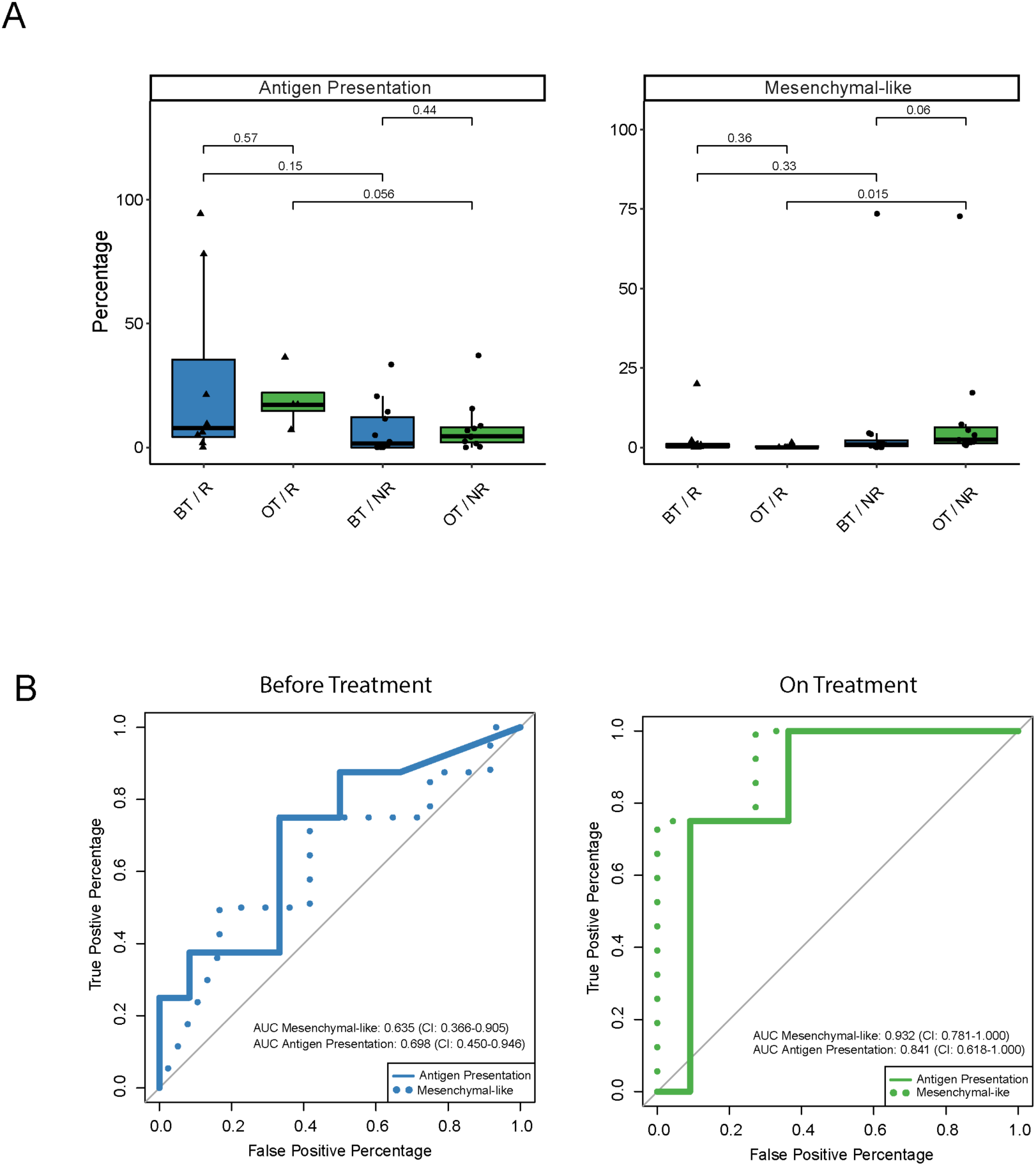
Antigen Presentation and MES states are associated with response to immunotherapy. **A,** Percentage of Antigen Presentation and MES cells out of all malignant cells in each sample, compared between R and NR at both time points (two-sided Wilcoxon test). **B,** Area Under the Receiver Operating Characteristics (AUROC) curves for the percentage of Antigen-presenting and MES cells out of all malignant cells in the BT (left) and OT (right) samples.

In contrast, the enrichment of melanoma MES cells in the NR lesions was only observed OT (Wilcoxon-test p=0.015). Importantly, the presence of both MES and Antigen Presentation cell populations in the OT samples showed a high diagnostic ability for response prediction and, thereby, biomarker potential (Figure 4B), whereas none of the other melanoma cell states showed any significant association with response (Supplemental Figure S7B).

### TCF4 orchestrates multiple melanoma transcriptional metaprograms

TCF4 is a known EMT inducer that promotes tumour progression and cell survival, in various epithelial cancers^51–54^. In melanoma, TCF4 was shown to promote invasion^55, 56^. In agreement with these observations, within the malignant compartment, *TCF4* is both specifically expressed and transcriptionally active in MES cells (Supplemental Figure S3A,B and Figure 3D). Consistent with these findings, *TCF4* expression was higher in the human TCGA samples harbouring the Verfaillie *et al.*^36^ invasive/mesenchymal-like (INV) compared to proliferative (PRO) melanoma signature, as well as in metastatic compared to primary lesions (Supplemental Figure S8A,B). *TCF4* expression also inversely correlated with *MITF* expression in samples from the TCGA cohort (Supplemental Figure S8C).

To assess the contribution of TCF4 in the establishment/maintenance of the MES transcriptional metaprogram, we performed bulk RNA-seq in a short-term melanoma MES line (MM099), following silencing of *TCF4* expression. Genes downregulated upon *TCF4* knockdown were involved in cellular movement, EMT, integrin signalling and angiogenesis, thus establishing its role as a driver of the MES transcriptional program (Figure 5A,B). This was concomitant to an upregulation of a series of MITF target genes and genes from the MEL transcriptional program (Figure 5B). This observation was consistent with a previous report indicating that TCF4 can repress MITF in normal melanocytes^57^. However, whether TCF4 represses MITF in melanoma cells is unknown. To test this hypothesis, we overexpressed *TCF4* in two different melanocytic melanoma cell lines (MM001 and MM011) and observed a downregulation of both *MITF* mRNA and protein levels as well as of its target genes (Figure 5C,D). These data indicated that, in addition to its function as a master regulator of the MES transcriptional program, TCF4 also actively suppresses the MITF-driven melanocytic transcriptional program. Importantly, silencing *TCF4* in MM099 caused a dramatic decrease in the ability to invade in short-term *in vitro* migration assays (Supplemental Figure S8D).

**Figure 5:**
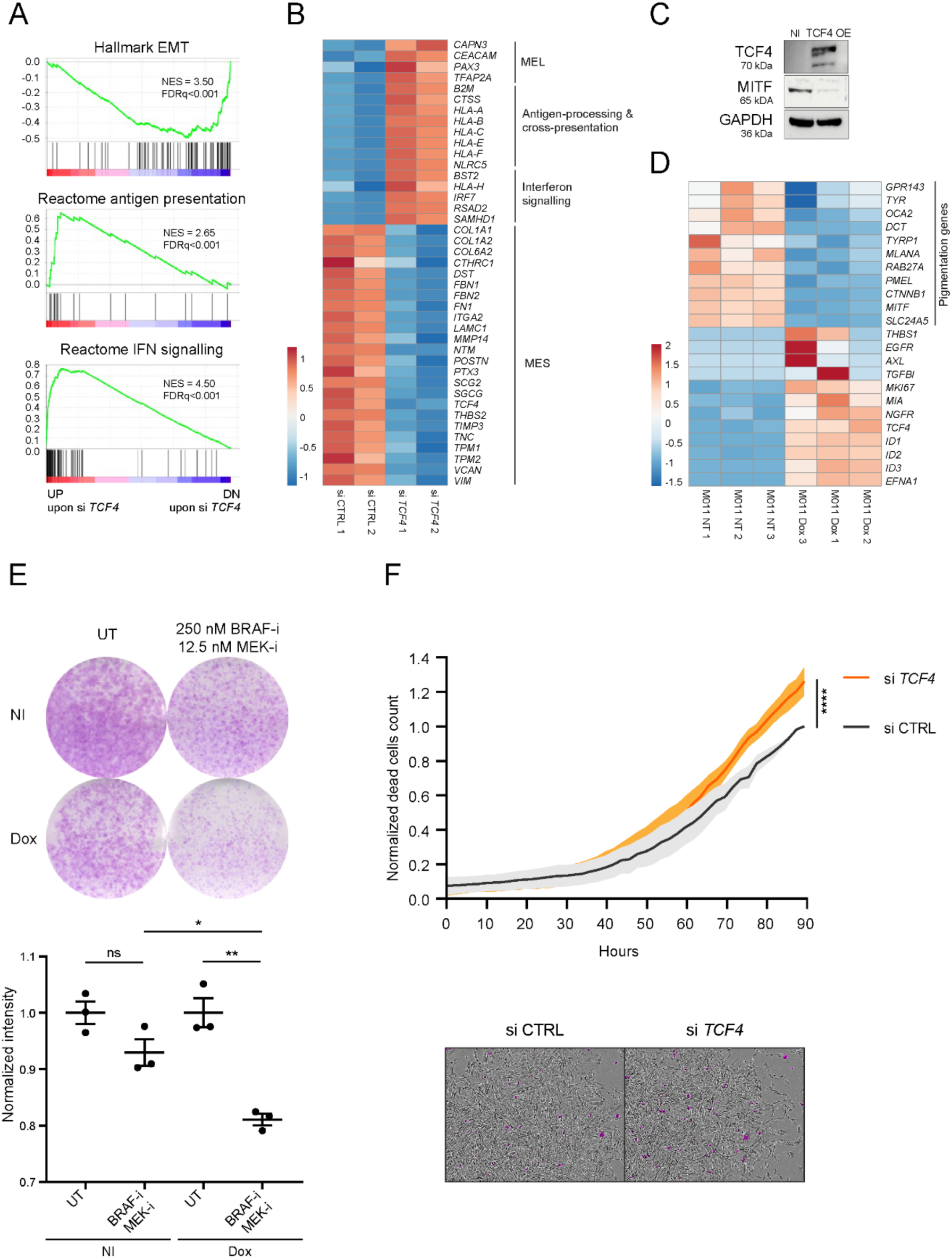
TCF4 orchestrates multiple melanoma transcriptional programs. **A,** GSEA analysis showing enrichment of gene sets related to EMT, antigen presentation, IFN signalling among the top DEGs upon silencing of *TCF4* in MM099 cells (bulk RNA-seq, n=2 biological replicates). NES, normalized enrichment score; FDR, false discovery rate. **B,** Corresponding heatmap of DEGs. **C,** Western blot analysis of TCF4, MITF and GAPDH expression in MM001 cells upon induction of TCF4 overexpression (OE). Expression in parental (non-induced, NI) cells is shown as control. **D,** Heatmap of differentially expressed genes in MM011 upon Doxycycline (Dox) -dependent induction of TCF4 expression (bulk RNA-seq, n=3 biological replicates). **E,** Colony formation assay performed with the melanoma MES cell line MM029, either left untreated (UT) or exposed to the BRAF- and MEK-inhibitors (Dabrafenib and Trametinib, respectively) at the indicated concentrations. TCF4 silencing was induced with Doxycycline (Dox). The non-induced (NI) cells are used here as control. Lower panel, quantification of ROIs (n=3 technical replicates), paired Student’s t-test, *p=0.0096, **p=0.0023). Error bars indicate mean ±SEM. **F,** Normalised dead cell counts in co-culture of MM099 with activated HLA-matched PBMCs upon silencing of *TCF4* by siRNA (n=3 biological replicates, paired t-test, ****p<0.0001). Line and filled area indicate mean ±SEM, respectively (n=6 technical replicates). Lower panel, representative IncuCyte images showing Caspase-3/7-positive cells in purple.

Consistent with the melanoma MES state being intrinsically resistant to MAPK therapeutics^11^, an inverse correlation between the sensitivity to BRAF- and MEK-inhibitors and *TCF4* expression was observed in the Cancer Cell Line Encyclopedia melanoma cell line cohort (Supplemental Figure S8E). Critically, silencing *TCF4* sensitized the human melanoma BRAF^V600E^-mutant invasive line MM099 to these inhibitors (Figure 5E). These data indicated that TCF4 contributes to the acquisition and/or maintenance of the mesenchymal-like phenotype and thereby to resistance to targeted therapy.

Remarkably, many genes involved in immune response (antigen processing and presentation, activation of leukocytes, and interferon signalling) were upregulated upon *TCF4* knockdown (Figure 5A,B). This included the transcription factor *NLRC5*, a master regulator of MHC class I and related genes^58^. Consistently, there was a strong enrichment of the Antigen Presentation and Interferon (IFN) signalling gene signatures among the genes upregulated upon *TCF4* silencing. Together, these data indicated that TCF4 actively suppresses the melanoma MEL, Antigen Presentation and Interferon signalling gene expression programs. By doing so, TCF4 may directly promote immune cell evasion and/or resistance to immunotherapy. Indeed, immune cells often target melanoma cells because they express melanocytic antigens. By suppressing the antigen processing and presentation machinery, TCF4 may further protect dedifferentiated melanoma cells from T-cell killing. Consistent with this model, *TCF4* silencing increased apoptotic cell death activation in a melanoma MES cell culture exposed to HLA-matched peripheral blood mononuclear cells (PBMCs), which were pre-treated with a T-cell activating cytokine cocktail (Figure 5F).

### Targeting TCF4 expression through BET inhibition

TCF4 was shown to drive B cell lymphoma and blastic plasmacytoid dendritic cell neoplasm (BPDCN)^54, 59^. In these studies, TCF4 expression was shown to be dependent on the bromodomain and extra terminal domain (BET) protein BRD4 through its recruitment to a specific TCF4 enhancer region. Inhibition of BRD4 using the BET-degrader ARV-771 was shown to decrease TCF4 expression. Interestingly, bulk Assay for Transposase-Accessible Chromatin using sequencing (ATAC-seq) revealed that the BRD4-binding enhancer region upstream from the *TCF4* promoter is largely accessible in melanoma MES cells such as MM099 and MM057, but not in melanoma MEL cells such as MM001 and MM011 (Figure 6A). Exposure of the melanoma line MM057 to the BET-degrader ARV-771 decreased chromatin accessibility upstream of the TCF4 locus, including of one of the sites previously identified as a BRD4-bound enhancer region (Figure 6A). Consistently, this treatment led to a dose-dependent decrease in *TCF4* expression (Figure 6B). Notably, the overall transcriptional reprogramming effect observed upon BET-inhibition was far more drastic in MES than in MEL cell lines (Figure 6C), indicating that the MES transcriptional program may be particularly dependent on BET epigenetic reader proteins.

**Figure 6:**
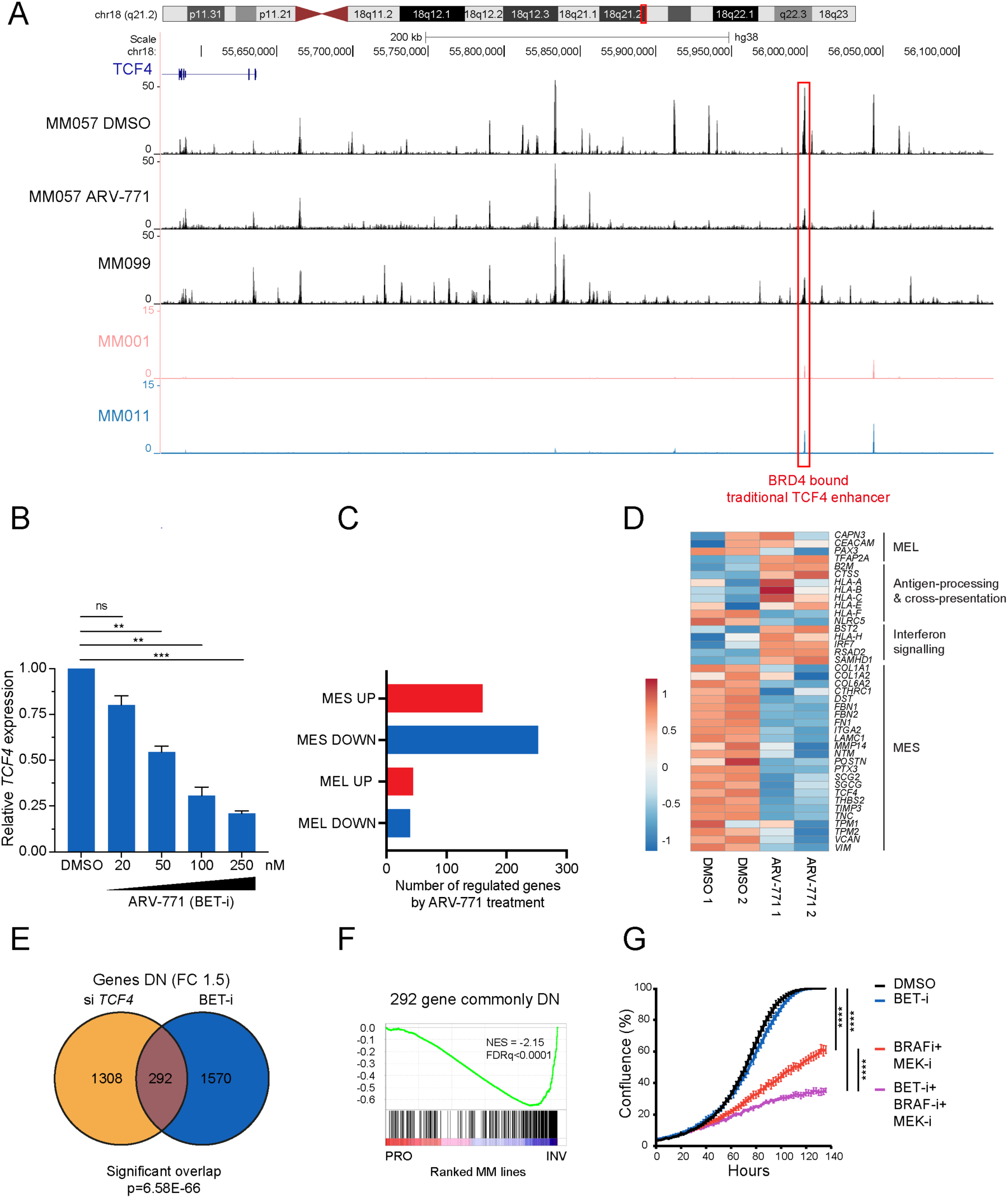
TCF4 targeting through BET-inhibition. **A,** ATAC-seq peaks indicating chromatin accessibility regions upstream the *TCF4* locus in MM057 control cells (DMSO) and following exposure to the BET-inhibitor ARV-771. The ATAC-seq profile of untreated MM099 and of two previously profiled^39^ melanocytic MM line (MM001, MM011) is shown below. The previously reported BRD4 binding site is framed^59^. **B,** RT-qPCR showing dose-dependent downregulation of *TCF4* upon ARV-771 treatment (n=3 technical replicates, Student’s t-test, **p<0.01, ***p<0.001). **C,** Number of up- or downregulated genes upon ARV-771 treatment in a panel of MM lines (bulk RNA-seq, n=2 biological replicates). MES, mesenchymal-like MM lines (MM047, MM099); MEL, melanocytic lines (MM034, MM164). **D,** Heatmap of selected panel of DEGs upon ARV-771 treatment in MM099 cells (bulk RNA-seq, n=2 biological replicates). **E,** Venn diagram showing the overlap between genes downregulated upon silencing of *TCF4* and ARV-771 treatment in MM099 cells (bulk RNA-seq, n=2 biological replicates, hypergeometric test) **F,** GSEA analysis showing enrichment of gene sets of a panel of proliferative versus invasive MM lines for the 292 commonly regulated genes. PRO, Proliferative; INV, Invasive. **G,** Cell growth of MM029 cells upon treatment with BET inhibitor (ARV-771 300 nM), BRAF- and MEK-inhibitors (Dabrafenib 50 nM, Trametinib 10 nM) or a combination thereof. Error bars indicate mean ±SEM (n=6 technical replicates, paired Student’s t-test, ****p<0.0001).

Furthermore, exposure to ARV-771 recapitulated most of the transcriptional changes observed upon TCF4 silencing (Figure 6D-F). Most genes from the MES signature were strongly downregulated, whereas genes from the MITF-dependent MEL signature were upregulated. Importantly, just like upon *TCF4* knockdown, these transcriptional changes were accompanied by an increased sensitivity to BRAF- and MEK-inhibition (Figure 6G). Moreover, the antigen presentation program was also upregulated upon exposure to ARV-771 (Figure 6D). This observation raises the possibility that this compound may also be used to increase the immunogenicity of MES cells and thereby their sensitivity to ICB.

## Discussion

In this study, we portrayed the cellular architecture of treatment-naïve skin and lymph node melanoma metastases, and thereby provide the field with a rich resource which may serve as the foundation for the creation of a comprehensive cell atlas of melanoma.

We focused our attention on the malignant compartment and show that the melanoma transcriptional landscape is even more complex than previously assumed. We describe a series of recurrent cell states that are evolutionarily conserved and show that the spatial distribution of these distinct melanoma subpopulations is not random, suggesting that the tumour microenvironment directly contributes to this geographically organised transcriptomic heterogeneity. This is, for instance, illustrated by the proximity we describe between the Antigen Presentation cell population and T cells, which suggests that this particular transcriptional program may be acquired through an intercellular communication pathway established by T cells. In concordance with this hypothesis, we previously observed that engraftment of a homogeneous mouse melanocytic cell line into immune competent, but not into immunodeficient, mice resulted in the formation melanoma lesions harbouring a complex and heterogenous transcriptomic landscape that included the Antigen Presentation cell state^32^ (data not shown). Moreover, a cancer cell state that expresses both antigen processing and interferon response genes was recently shown to recur across multiple tumour types and to colocalize with T cells^60^.

The functional contribution to tumour growth and/or metastatic spreading of these newly defined melanoma transcriptional states remain to be explored. The identification of these evolutionarily conserved states makes it possible to use the mouse as a model system for this, through lineage tracing and depletion experiments. Notably, using such approaches, we recently demonstrated that a population of MES cells present in primary tumours in minute amounts drives the metastatic process^32^.

Importantly, our single-cell RNA-sequencing data made it possible to identify true malignant MES cells and develop a set of markers that unambiguously distinguish MES cells from CAFs in scRNA-seq datasets, as well as *in situ*. The method we describe herein is a critical step forward for the field, as it permits to directly assess the contribution of this critical cell population to various aspects of melanoma biology and map their position within the complex melanoma ecosystem. In addition, the gene signature established from our single-cell datasets may also, in theory, be used to infer the proportion of MES cells from bulk transcriptomics data. However, although we have not tested the performance of our MES signature to predict the presence of these cells in bulk RNA-sequenced samples, we anticipate that, given the overall rarity of these cells in our melanoma samples, obtaining reliable deconvolution results may be very challenging. Such deconvolution analyses should therefore be performed with extreme caution.

Previous studies indicated that melanoma de-differentiation may contribute to immune escape^12, 13^. However, a clear association between melanoma MES and resistance to immune checkpoint inhibition is yet to be formally established. Our data provide evidence that these cells may contribute to primary resistance to ICB therapy and that their presence after one cycle of treatment is predictive for a lack of response. This observation indicates that (multiplex) analysis of an early on-treatment biopsy (2 weeks after the first infusion of immune checkpoint inhibitors) may provide a predictive biomarker for robust stratification of patients into R and NR, and thus before NR patients are increasing their risk of developing treatment related adverse events. This observation validates our initial hypothesis that early on-treatment samples may be much more informative than baseline samples. It is, however, important to stress that the predictive value of the presence of MES cells needs to be firmly established in a larger population cohort before this concept can be exploited clinically.

Although the presence of MES in the on-treatment samples is predictive of lack of response, their proportion remains overall relatively low (below 20% of all malignant cells for most samples) at this early time point. Additional studies will be needed to monitor the dynamics of this population over time and assess whether their proportion increases at later time points. However, this observation also suggests that these cells are not the only melanoma cells able to escape T cell killing. One interesting possibility is that the MES population may also contribute to primary resistance to ICB in a non-cell autonomous manner, by promoting an immunosuppressive environment. In support of this possibility, emerging data indicate that cells harbouring overlapping phenotypes with melanoma MES cells, such as inflammatory fibroblasts and mesenchymal carcinoma cells, do secrete immunosuppressive factors such as CD73^61^. Moreover, it was shown that expression of the EMT TF ZEB1 in melanoma cells is associated with decreased CD8^+^ T cell infiltration and ZEB1 ectopic expression in melanoma cells impairs CD8^+^ T cell recruitment in syngeneic mouse models, resulting in tumour immune evasion and resistance to ICB therapy. Mechanistically, it was shown that ZEB1 directly represses the secretion of T cell-attracting chemokines, including CXCL10^62^.

We identified TCF4 (E2-2) as a key driver of the mesenchymal-like transcriptional program. Other TFs such as c-JUN/AP1^63^, ZEB1^62^, SOX9^64^, TEADs^36^ and more recently PRRX1^32^. have previously been identified as master regulators of this particular program. It will be thought-provoking to study how redundant and interconnected the activity of these TFs are.

Critically, our data suggest that in addition to driving the MES transcriptional program and related phenotypes, TCF4 actively suppresses both the MEL and antigen presentation programs. By doing so, TCF4 may directly promote immune cell evasion and resistance to ICB therapy as melanocytic antigens are prime targets of the adaptive immune system. Moreover, by suppressing the antigen processing and presentation machinery, TCF4 may further reduce the immunogenicity of this dedifferentiated melanoma subpopulation. Together, these data identify TCF4 as a putative target to improve response to ICB therapy.

A potential limitation of targeting TCF4 is that this TF is also expressed in other cell types. However, beside melanoma MES cells, TCF4 expression is the highest in pDCs, where it was shown to act as a major suppressor of their immunogenic function^65^. Therefore, manipulation of the TCF4 pathway in pDCs could represent a therapeutic opportunity to further boost antitumor immunity.

Additionally, TFs are notoriously difficult to target pharmacologically. However, just like observed in BPDCN cells^54, 59^, BET-inhibition recapitulated most of the transcriptional changes observed upon TCF4 silencing in melanoma MES cells. The use of BET protein inhibitors may therefore offer an alternative strategy to target this pathway. Mechanistically, the disruption of the TCF4-controlled transcriptional program by BET inhibitors can be explained either by the dependency of TCF4 expression itself on the recruitment of BRD4 to the TCF4 promoter and/or by an important role of BDR4 in the recruitment of TCF4 to, at least some of, its target genes. Additional experiments will be required to further establish the TCF4-dependency of the effects observed upon BET-inhibition in melanoma MES cells and to discriminate between these two, not necessarily mutually exclusive, scenarios.

The treatment of cancer with BET-inhibitors has been explored in early clinical trials^66^. Toxicity profiles of several generations of inhibitors showed that these agents can be given safely to patients. Unfortunately, these inhibitors have not yet been broadly used in the clinic due to their modest anti-tumour activity when used as single agents. BET-inhibitors remain, however, attractive drugs for combinatorial treatments and when used in the appropriate clinical settings^67, 68^. Our observation that BET-inhibition sensitizes melanoma cells to BRAF- and MEK-inhibition, is in line with similar observations by other groups^69–71^, and offers one clinical context in which BET-inhibitors may provide clinical benefit for BRAF-mutant melanoma patients (Supplemental Figure S9). Another attractive clinical context in which BET-inhibition could be positioned is in combination with ICB. We provide evidence that exposure of melanoma MES cells to the BET-inhibitor ARV-711, just like TCF4 silencing, unleashes the expression of antigen presentation machinery and HLA-genes. These data therefore offer a rationale to increase the immunogenicity of melanoma MES cells and warrant the further testing of BET-inhibition in combination with ICB to overcome primary resistance (Supplemental Figure S9). Notably, recent preclinical studies have supported this possibility^69, 72, 73^. Moreover, since emergence of the mesenchymal-like signature was shown to be prominent in patients who experience disease progression after first line immunotherapy^17^, one could envision that BET-inhibition could reinvigorate anti-tumour immune responses and overcome secondary resistance to ICB (Supplemental Figure S9).

Lower efficacy was observed with ICB therapy when given as second-line treatment, after first-line targeted^74–76^. It has recently been proposed that this cross-resistance phenomenon may be driven, at least partly, by changes in the tumour microenvironment induced by BRAF and MEK-inhibition, leading to a lack of functional CD103^+^ DCs, and consequently an ineffective T cell response^77^. Our findings may offer an alternative (but not mutually exclusive) explanation, invoking a cancer-cell intrinsic mechanism. It is well-established that melanoma MES cells are key drivers of tolerance and/or resistance to targeted therapy^11^. Likewise, we show that this population is enriched in (early on-treatment) lesions from non-responders to ICB, and therefore propose that MES cells may drive, at least partly, cross-resistance to these treatments. Importantly, we show that these cells are exquisitely sensitive to the BRAF/MEK/BET-inhibitors triple combination. This combination may therefore also offer an attractive treatment strategy for patients who do not respond to immunotherapy and those who develop resistance to targeted therapy through nongenetic mechanisms (Supplemental Figure S9).

Together, our data offer the rationale for the (pre-)clinical testing of BET-inhibition (or TCF4 targeting) as both a putative sensitizer to targeted therapy and ICB and for the treatment of patients that develop secondary resistance to these therapies. We argue, however, that the testing of these new combination treatment regimens should be accompanied by a careful selection of the models and patients. In this context, the method we describe herein, which allows for the unambiguous identification of melanoma MES cells in tumour biopsies, should be considered as a critical selecting or recruiting criteria.

## Acknowledgments

We thank Prof. G. Ghanem, LOCE-Institut J. Bordet, Université Libre de Bruxelles for providing us with the MM lines. We thank Odessa Van Goethem and Veronique Benne for their assistance with the experiments. We thank Aurélie Bousard for the analysis of the OmniATAC-seq experiment. P.K. received financial support from Marie Curie Individual Fellowship (H2020-MSCA-IF-2018, #841092). J.P. received financial support from Marie Curie Individual Fellowship (H2020-MSCA-IF-2019, #896897). S.C. received a PhD fellowship from FWO (#1SD1620N). D.P. received a PhD fellowship from the VIB PhD international program. A.N. received postdoctoral fellowship from the KU Leuven (PDMT1/21/035). F.R. acknowledges support from the Alexander von Humboldt Foundation. S.K, K.V and T.V. are supported by KU Leuven (Opening The Future, SymBioSys - C14/18/092, IDN/19/039; IDN/21/006), the Research Foundation Flanders (FWO, grant I001818N). A.A. was partially funded by the KU Leuven funding (C3/19/053), Opening the Future Campaign of the KU Leuven University Fund and by Kom op tegen Kanker (Stand up to Cancer), the Flemish cancer society. F.M.B. was funded by KUL INTERNE FONDSEN MIDDEL-Zware infrastructuren EMH-D8191-AKUL/19/30 I005920N, FWO Fundamenteel Klinisch Mandaat EMH-D8972-FKM/20, Impulsbudget UZ Leuven 2018.The computational resources and services used in this work were provided by the VSC (Flemish Supercomputer Centre), funded by the Research Foundation - Flanders (FWO) and the Flemish Government – department EWI. This work has received funding within the Grand Challenges Program of VIB and from FWO (#G0C530N and G070622N), Stichting tegen kanker (FAF-F/2018/1265), Neftkens foundation, KULeuven (C1 grant) and the Belgian Excellence of Science (EOS) program to J-C.M. W.A. received financial support from C14/21/095-InterAction and E.L. received from the Melanoma Research Alliance (https://doi.org/10.48050/pc.gr.80542) and FWO-KOTK grant (#G097620N). The schematic figure of the study was created with BioRender.com.

## Author contributions

J.P. analysed and interpreted the human scRNA-seq and untargeted spatial transcriptomic data. F.R. analysed and interpreted TCGA data, and performed TCF4 related RNA-seq and invasion assays. D.P. conducted most cell culture experiments, acquired, analysed and interpreted the data, being partly supervised by F.R and W.A. G.B. and E.L. performed all scRNA-seq experiments. D.L. coordinated the generation of the scRNA-seq data. E.L. and Y.v.H. curated patient clinical data. E.L. analysed Akoya/RNAscope data. Y.v.H performed the MILAN experiments. A.A.M., F.B. and Y.v.H analysed and interpreted the MILAN data. A.N. contributed cell culture experiments and generation of cell lines. P.K. and F.V. contributed to the analysis of the CODEX and mFISH data. S.M, L.V, contributed to the generation of CODEX and mFISH data. E.L., S.M, S.K, K.V. and T.V. contributed to the generation of spatial transcriptomics (Visium, 10X Genomics) data. S.C. and E.Le. developed and supervised the HLA-matched PBMC killing assay. V.B. collected the clinical biopsies. J.vd.O. and F.B. provided pathology support. F.R., O.B, J-C.M conceptualized, designed the clinical and/or research study. J-C.M and J.P. wrote the manuscript. All authors read and edited the manuscript.

## Methods

### Patient biopsies

Tumour biopsies were collected as part of a non-interventional prospective study investigating transcriptomic changes upon immune checkpoint inhibition (Prospective Serial biopsy collection before and during immune-checkpoint inhibitor therapy in patients with malignant melanoma; SPECIAL). While most patients (n=20) were treatment-naïve, patients with metastatic relapse were allowed prior systemic treatment in adjuvant setting (n=2). In addition, one patient had received Cisplatinum-based neo-adjuvant chemotherapy for a metachronous non-small cell lung carcinoma eight months before inclusion. Written informed consent was obtained from all patients. All study procedures were in accordance with the principles of the Declaration of Helsinki, applicable Belgian law and regulations, and approved by the UZ Leuven Medical Ethical Committee (S62275).

### Tumour dissociation of human samples

Fresh tumour tissue was collected in cold transport Dulbecco’s Modified Eagle Medium (DMEM, Invitrogen, Cat#61965025) on ice. To make a single cell solution, tumour fragments were rinsed in cold Dulbecco’s Phospho-Buffered Saline (DPBS) and mechanically and enzymatically dissociated. The tumour was minced with sterile scalpels and incubated 15 minutes in a heather-shaker at 37°C 800 rpm in 1,32 mL DMEM supplemented with 120 µL DNase I (10 mg/mL; Sigma-Aldrich, Cat#11284932001) and 60 µL collagenase P (50 ng/mL; Sigma-Aldrich, Cat#11249002001). The sample was diluted 1:2 in DPBS, centrifuged 5 min. at 300G at room temperature, incubated 5 min. in 500 µL red blood lysis buffer at room temperature, washed twice with DPBS supplemented with 0,04% bovine serum albumin and strained through a 35 µm nylon mesh. Cell concentration and viability was determined with acridine orange/propidium iodide staining (Westburg, Cat#LB F23001) on a LUNA-FL automated fluorescence counter.

### Single-cell RNA-sequencing

Libraries for scRNA-seq were constructed using the 10X Genomics Chromium platform according to manufacturer’s instructions. Library construction was primarily done with the Chromium Single Cell 3ʹ GEM, Library & Gel Bead Kit v3 (10x genomics, Cat#1000092). Thirteen samples were processed using the Chromium Single Cell A Chip Kit and 5’ Library & Gel Bead Kit (10x genomics, Cat#1000014). When comparing sequenced 3’ and 5’ gene expression libraries from the same tumour samples, we observed similar quality metrics. We opted for high target recovery (median 5000, range 1000-10000), keeping within the range of optimal input concentration per target recovery, as recommended by the manufacturer. In brief, cells were partitioned into Gel Bead-in-emulsions (GEMs) at limiting dilution, where lysis and reverse transcription occurred yielding uniquely barcoded full-length cDNA from poly-adenylated mRNA. GEMs were subsequently broken, and the pooled fraction was amplified, followed by fragmentation, end repair and adaptor ligation of size selected fractions.

All libraries were sequenced with single end reads on an Illumina NextSeq, HiSeq4000 or NovaSeq6000 until sufficient saturation was reached (60% on average). The raw sequencing reads were processed by CellRanger (10x Genomics), human reference genome v. GRCh38.

### scRNA-seq data analysis

Raw count matrices were analysed using R package Seurat v. 3.1.5^78^. The matrices were filtered by removing cell barcodes with >1000 expressed genes, <7,500 expressed genes and <30% of reads mapping to mitochondrial reads. Next, SCTransform was applied to each Seurat object for data normalization and transformation. DoubletFinder v. 2.0.283 was applied to each Seurat object (sample) separately assuming that the doublet rate in each sample was as indicated in the 10X Genomics website. Next, all the Seurat objects were merged, SCTransform was applied regressing out mitochondrial read percentage per cell. Subsequently, the data integration was performed using R package Harmony v. 1.040. After having the data normalized and integrated, cell cycle scoring was performed, data were filtered for singlets, and SCTransform was applied regressing out mitochondrial read percentage and cell cycle scores. This was followed by data integration of this subset as described above. The number of dimensions for clustering were chosen based on Harmony embeddings clustering. The cut-off was driven by identification of clear variation in embeddings across the cells. For cell type identification from the malignant, immune and stromal compartments we analysed the data including cells from both time points, but for the detailed characterisation of the treatment naïve samples we subset only for this time point.

### Initial identification of the tumour microenvironment compartments

To gain a global view on the components of the tumour microenvironment we used existing signatures acquired from Jerby-Arnon *et al.*^22^ to calculate gene set scores using R package AUCell v.1.6.1^31^ for the immune, stromal and malignant compartments. By plotting the scores, we assigned each unsupervised cluster to one of these three compartments.

### CNV inference in human samples

To distinguish malignant from normal cells we inferred copy number variation (CNV) based on scRNA-seq data using the R package HoneyBadger v. 0.1^79^. The count matrix from the “RNA” assay of the integrated Seurat object of all cells was used as input. Immune cells were used as a reference for normal cells. They were defined based on the immune gene set from Jerby-Arnon et al.^22^, using an AUCell score cut-off >0.15.The mean CNV score was calculated as below:

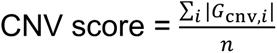

where, G = gene, *i* = cell.

### Identification and analysis malignant cells

Differential gene expression was run between globally classified malignant clusters (2,28,0,12,17,20,19) vs CAFs clusters (7,8) (Supplemental Figure S1A) using Seurat FindMarkers function (two-sided Wilcoxon test). Next, for each gene, the difference of the percentage of cells expressing this gene in the malignant clusters minus the CAFs clusters was calculated and the genes were sorted in descending order. The top 50 genes were plotted on the global UMAP in order to identify the most specific and ubiquitously expressed ones within malignant clusters further called as Melanoma Score (MS).

To identify malignant cells, three stringent steps of filtration were applied. Firstly, the data was subset based on the AUCell score of malignant gene set acquired from Jerby-Arnon *et al.*^22^ >0.11 or mean CNV score >0.15. Subsequently, cells that passed the first filtration step were filtered based on the MS >0.2 or mean CNV score >0.15. Finally, to remove any contaminating immune cells, we filtered out *PTPRC* (CD45) cells. Lastly, samples with less than 10 remaining cells were removed from downstream analysis.

The malignant cell subsets were subjected to SCTransform (regressing out mitochondrial read percentage and the cell cycle scores) and Harmony integration (grouping the variables by samples) followed by unsupervised Seurat clustering. The number of dimensions for clustering were chosen based on Harmony embeddings clustering – a cut-off was driven by identification of clear variation in embeddings across the cells. Number of clusters was chosen based on Silhouette^29^ scores measured at different resolutions and biological relevance of the marker genes per cluster. The marker genes of each unsupervised and semi-supervised cluster were identified using FindAllMarkers function in Seurat (two-sided Wilcoxon test). The final cluster annotations were based on the enriched pathways and terms of the top marker genes per cluster (top 100 genes) using an online tool Enrichr (https://maayanlab.cloud/Enrichr/). To understand the biological identity of the malignant clusters, we used databases such as Gene Ontology (GO), Reactome, OMIM Disease, MSigDB Hallmark 2020, Jensen Compartments, CellMarker Augmented, and CCLE Proteomics. Furthermore, AUCell scores of the functionally enriched marker genes per malignant mouse state (acquired from Karras *et. al*^32^), the top 100 marker genes of the malignant states, and the scores of various previously published melanoma signatures were averaged and plotted across the malignant clusters.

To infer cell-cell interactions we used the Seurat object and run CellChat^80^ version 1.1.3 applying 10 % truncated mean for average gene expression per cell group and minimum twenty cells required per cell.

The percentages of the malignant clusters within the malignant compartment were calculated per sample and tested among various groups. For the two groups comparisons, the two-sided Wilcoxon test, for three or more groups, the Kruskal-Wallis test was used. Area Under the Receiver Operating Characteristics (AUROC) was used to estimate the response prediction.

### Gene regulatory network analysis

SCENIC^31^ analysis was run with raw counts from the “SCT” assay of malignant cells 50x. SCENIC uses gene regulatory network inference, followed by a refinement step using cis-regulatory information, to generate a set of refined regulons (i.e. TFs and their target genes) in the scRNA-seq data. The Python implementation, (pySCENIC: https://github.com/aertslab/pySCENIC, version 0.9.19), was run using a Nextflow pipeline (https://github.com/aertslab/SCENICprotocol, version 0.2.0), which streamlined the main steps of gene regulatory network inference and refinement with pySCENIC, as well as the quantification of cellular activity, and visualization. The Nextflow pipeline also performed a standard analysis in parallel, using highly variable genes selected based on expression. Differentially activated TF regulons of each malignant cluster were identified by the two-sided Wilcoxon test (using Bonferroni correction for multiple tests) against all the cells of the rest of the clusters.

### Identification of the Minimal Lineage Gene signature (MLGs)

To identify the MLGs we subset CAFs together with the Mesenchymal-like state and performed differential gene expression analysis between them. The top 50 genes were called the MLGs, from which four (*SOX10*, *S100A1*, *MITF* and *CDH19*), were selected for further validation by CODEX/mFISH (RNAscope).

### Validation of identified melanoma states in independent scRNA-seq dataset

Transcript per Million (TPM) normalized the Jerby-Arnon *et al.*^22^ dataset was downloaded from the GEO portal (Accession number GSE115978). The TPM normalized dataset was used to generate a Seurat object. The cells with a number of genes > 1000 & < 7500 were selected for further analysis. Next, the AUCell score for the MS was calculated using the same set of genes as in the main cohort (*EDNRB*, *MYO10*, *PLP1*, *ERBB3*, *SYNGR1*). The malignant cells were subset based on an SMS score >0.1 and the criteria applied in the Jerby-Arnon *et al.*^22^ study. Additionally, cells positive for *PTPRC* were excluded. Next, the data was scaled regressing out the cell cycle scores and percentage of expressed mitochondrial genes, and integrated using Harmony.

To validate the transcriptomic states identified in our dataset we performed label transfer of the malignant clusters in Seurat, using the following parameters: integration features = 3000, k.anchor = 20, and reduction = “pcaproject”. The prediction scores were plotted on the Harmony integrated UMAP.

### Spatial transcriptomics

Selected samples were processed for spatial transcriptomics using the 10X Genomics Visium platform. The analysis of these was approved by the Ethical Commission of the University Hospital of Leuven and approved by the review board (#S55760). Tumours were dissected, washed with 1x DPBS and snap-frozen in liquid nitrogen-chilled isopentane. Frozen tumours were transferred to a cold tissue mould filled with chilled optimal cutting temperature compound (Tissue-Tek O.C.T. compound, Sakura Finetek Cat#4583). The mould was then immediately placed on dry ice. Tissue blocks were stored at −80°C in a sealed container. Both the tissue block and the proprietary Visium Spatial Gene Expression Slide (10X Genomics, Cat#PN-2000233) were equilibrated inside the cryostat for 30 min at −12 °C before sectioning. Sections were cut at a thickness of 10 µm and immediately placed onto the slide. Slides containing sections were stored at −80°C for a maximum of 24h before use.

Fixation, staining, imaging, and construction of cDNA libraries was done according to the manufacturer’s instructions (Visium Spatial Gene Expression User Guide_Rev D; 10x Genomics, CG000239) using the Visium Spatial Gene Expression Slide & Reagent Kit (10x Genomics, Cat#PN-1000187). Briefly, sections were fixed in chilled methanol for 30 min at −20 °C and stained with haematoxylin and eosin. Imaging was performed on a Nikon-Marzhauser Slide Express 2 whole-slide scanner at 10x magnification. After imaging, sections were permeabilized at 37 °C for 18 minutes. Permeabilization time was determined using the Visium Spatial Tissue Optimization Slide & Reagent Kit (10x Genomics, PN-1000193) following the Visium Spatial Tissue Optimization User Guide_RevA (10x Genomics, CG000238). After permeabilization, the on-slide reverse transcription reaction was performed at 53 °C for 45 min. Second strand synthesis was subsequently performed on-slide for 15 min at 65 °C. All on-slide reactions were performed in a thermocycler with a metal slide adapter plate. Following second strand synthesis, samples were transferred to tubes for cDNA amplification and clean-up. Library QC was assessed using an Agilent Technologies Bioanalyzer High Sensitivity kit (Agilent Technologies, Cat#5067-4626).

Visium libraries were sequenced on Illumina NextSeq2000. The sequencing depth was chosen by determining the amount of 55µm spots that were covered by tissue, and this was multiplied by 50.000 reads. Raw sequencing files were processed with SpaceRanger (v1.1.0, 10x Genomics) to generate spatial gene expression matrices. Next, the data was analysed using the Seurat v. 4.1.0^81^ spatial vignette in R v. 4.0.2. Spots with spatial features >500 and percentage of mitochondrial reads either <3 or <5 were retained and the expression data was normalized using SCTransform. Firstly, the spatial distribution of the major tumour immune microenvironment (TIME) constituents such as malignant cells, T cells, B cells, macrophages, CAFs, and ECs was mapped using the label transfer function with CCA-based label transfer (k anchor=10). Furthermore, to identify true malignant spots, we leveraged copy number inference using HoneyBadger^79^. Bcell or EC spots with prediction score >0.8 were used as “normal” reference. Subsequently, we selected spots with a prediction score >0.7 for the malignant label and the mean CNV score (calculated as described above) >0.07 and annotated these as melanoma. Finally, for the malignant cell deconvolution, distance and co-localization calculations we used the CellTrek^43^ R package.

### Multiplex immunostaining followed by multiplex FISH

Five µm FFPE tissue sections of selected samples sectioned 5 were cut and mounted on poly-L-lysine coated coverslips. Akoya Biosciences CODEX multiplex immunostaining (CODEX) and ACDbio RNAscope HiPlex v2 (12-plex) multiplex FISH (RNAscope) were each performed according to their respective manufacturer’s instructions (kits used are listed in Supplemental Table S8), and combined in sequence as previously described, with slight modifications^82^.

In brief, coverslips were deparaffinized followed by heat-induced antigen retrieval in citrate buffer, pH 6. Next, they were stained with a combination of DNA-barcoded primary antibodies, including in-house conjugated antibodies (Supplemental Table S8), washed and post-fixed in ice-cold methanol. They were mounted on an Akoya Biosciences CODEX system for multiple cycle immunostaining and imaged using a Keyence microscope with Akoya Biosciences CODEX instrument manager and Keyence software. Secondary antibodies were fed to the instrument in a pre-prepared 96-well plate. In total, 11 cycles of immunostaining (including 2 blanks with only nuclear staining) were run, consisting of DAPI nuclear staining, Atto550-, Cy5- and Alexa Fluor 750 fluorophores. Akoya Biosciences CODEX processor software performed automated image registration, autofluorescence and background subtraction. Cover slips were kept in storage buffer until RNAscope was performed. To prepare samples for RNAscope, samples were washed in ethanol for 2 min. and air dried for 5 min. in a 60°C oven. Target retrieval was followed by protease treatment. The 12-plex RNAscope assay consisted of 3 rounds (each round using 4 probes) of probe hybridization, amplification, autofluorescence reduction, fluorophore hybridization, DAPI counterstaining, imaging, fluorophore cleavage and washing. Coverslips were imaged using VectraPolaris Automated Quantitative Pathology Imaging System.

CODEX images were registered to RNAscope using the BigWarp plugin for ImageJ^83^ The CODEX image stack was used as a fixed target to register the 3 RNAscope imaging rounds onto, using manually placed landmarks. This resulted in resampling of the RNAscope to target resolution. Next, regions of interest (ROIs) of 100 x 100 µm were delineated using QuPath Quantitative Pathology & Bioimage Analysis software^84^. In these ROIs, the autofluorescence channel of each RNAscope imaging round was subtracted from each respective fluorescent channel using the Image Calculator in ImageJ. Cells were segmented with the StarDist^85^ extension in QuPath, using the dsb2018_heavy_augment.pb pretrained model^86^.

### MILAN (mIHC)

Multiplex immunofluorescent staining was performed according to the previously published MILAN protocol^44^. Immunofluorescence images were scanned using the Axio scan.Z1 slidescanner (Zeiss, Germany) at 10X objective with resolution of 0.65 μm/pixel. All samples were stained simultaneously. Image acquisition order was distributed spatially and independently of patient replicates. The stains were visually evaluated for quality by digital image experts and experienced pathologists (FB, YVH, double-blind). Multiple approaches were taken to ensure data. On the image level, focus, presence of external artefacts and tissue integrity were reviewed. Regions that contained severely deformed tissues and artefacts were identified and excluded from downstream analysis. Antibodies that gave low confidence staining patterns by visual evaluation were excluded from the analysis. Image analysis was performed following a custom pipeline. Briefly, flat field correction was performed using a custom implementation of a previously described algorithm^87^. Then, adjacent tiles were stitched by minimizing the Frobenius distance of the overlapping regions. Next, images from consecutive rounds were registered following an algorithm previously described^88^. During this process, the first round was always used as a fixed image whereas all consecutive rounds were sequentially used as moving images. Transformation matrices were calculated using the DAPI channel and then applied to the rest of the channels. Registration results were visually inspected by domain experts (FB, YVH). Samples with tissue folds showed significant misalignments and were manually segmented in different regions. Each region was independently re-registered. Downstream analysis was independently performed for each annotated region. Next, tissue autofluorescence was subtracted using a baseline image with only secondary antibody. Finally, cell segmentation was applied to the DAPI channel using StarDist^85^. For every cell, topological features (X/Y coordinates), morphological features (nuclear size), and molecular features (Mean Fluorescence Intensity (MFI) of each measured marker) were extracted.

For the cell Identification MFI values were normalized within each region to Z-scores as recommended in Caicedo et al^86^. Z scores were trimmed in the [0, 5] range to avoid a strong influence of possible outliers in downstream analyses. Single cells were mapped to known cell phenotypes using three different clustering methods: PhenoGraph^89^, FlowSom^90^, and KMeans as implemented in the Rphenograph, FlowSOM, and stats R packages. While FlowSom and KMeans require the number of clusters as input, PhenoGraph can be executed by defining exclusively the number of nearest neighbours to calculate the Jaccard coefficient. The number of clusters identified by PhenoGraph was then passed as an argument for FlowSom and KMeans. Clustering was performed exclusively in a subset of the identified cells (50,000) selected by stratified proportional random sampling and using only the 23 markers defined as phenotypic. For each clustering method, clusters were mapped to known cell phenotypes following manual annotation from domain experts (FMB, YVH, double-blind). If two or more clustering methods agreed on the assigned phenotype, the cell was annotated as such. If all three clustering methods disagreed on the assigned phenotype, the cell was annotated as “not otherwise specified” (NOS). Annotated cells were used to construct a template that was in turn used to extrapolate the cell labels to the rest of the dataset. To that end, a UMAP was built by sampling 500 cells for each identified cell type in the consensus clustering. The complete dataset was projected into the UMAP using the base predict R function. For each cell, the label of the closest 100 neighbours was evaluated in the UMAP space and the label of the most frequent cell type was assigned.

Melanoma cells were further segmented based on the expression of HLA-DR^91^. Here, we set a cut-off of Z=2 to differentiate between HLA-DR positive and HLA-DR negative melanoma cells.

For neighbourhood analysis, a quantitative analysis of cell-cell interactions was performed using an adaptation of the algorithm described in Schapiro, *et al.*^92^. A detailed description of the adapted implementation was previously published^93^. Briefly, for every cell, all the other cells that are located at a maximum distance d were counted. Then the tissue is randomized preserving the cytometry of the tissue as well as the X and Y coordinates of each cell but permutating the cell identities. This is repeated N times (here N=1000) which allows to assign an empirical p-value by comparing the number of counts observed in the real tissue versus the N random cases. We performed the described analysis for different values of the distance d (from 10 to 100 µm with a step of 10 µm) to show the consistency of the reported results. Particularly here, the analysis was performed exclusively to evaluate whether CD3+ and CD8+ T cells (Tcys) were interacting more with HLA-DR positive or HLA-DR negative melanoma cells. Therefore, we only included melanoma subtypes in the randomization process while keeping all the other cell subtypes unchanged. To add an effect-size metric, we also calculated the ratio between the observed counts and the random counts.

### Cell culture

The human melanoma cell cultures were derived from patient biopsies by the Laboratory of Oncology and Experimental Surgery (Prof. Dr. Ghanem Ghanem, Institute Jules Bordet, Brussels, Belgium). All cell lines (MM011, MM029, MM034, MM047, MM057, MM099, MM164) were grown in 5% CO2 at 37°C in F10 supplemented with 10% FBS, 2.5% GlutaMAX and 1% penicillin/streptomycin. HEK293 FT cells were grown in DMEM with 10% FBS and 1% penicillin/streptomycin. Cells were tested for Mycoplasma contamination prior to performed experiments.

### Drugs

Dabrafenib (Cat#HY-14660) and Trametinib (Cat#HY-10999) were purchased from MedChemExpress. ARV-771 (Cat#HY-100972) was purchased from Bioconnect.

### siRNA-Mediated Transient Genetic Inactivation

Cells were transfected with the indicated specific short interfering RNA (siRNA) SMARTpools (Dharmacon, Cat# L-004594-00-0005 and Cat# D-001810-10-20) using TransIT-X2 Transfection Reagent (Mirus) according to the manufacturer’s protocol. siRNAs were used at a final concentration of 50 nM.

### Lentiviral vector production

HEK293 FT cells were transfected with dVPR and VSVG packaging plasmids using Lipofectamine 2000 reagent (ThermoFisher Scientific) according to the manufacturer’s instructions. 24 hours after transfection, medium was replaced with DMEM medium (Invitrogen) supplemented with 20% foetal bovine serum (FBS). Medium containing viral vectors was collected 48 and 72 hours after transfection. Viral vectors were filtered through a 0,45 nm syringe filter, aliquoted and stored at −80 °C.

Inducible TCF4 overexpression was achieved using a Doxycycline-inducible vector system. Briefly, a TetR-T2A-NeoR insert was cloned inside a FUGW vector (Addgene Plasmid, Cat#14883). In a second plasmid, the expression of TCF4 was controlled by Doxycycline through a TetO-regulated CMV promoter. Both vectors were transduced in MM011 cells and selection was obtained with neomycin and puromycin respectively. Inducible TCF4 downregulation was achieved using a Doxycycline-inducible vector system. Briefly, the shRNA sequence targeting TCF4 was designed based on the sequence of the siRNA pool and cloned in a FH1 vector (Addgene Plasmid, Cat#164098).

### Bulk RNA-sequencing

Approximately 2*10^5^ cells were plated in a 6-well plate. For knockdown experiments, these were transfected with the described siRNA pool 24 and 72 hours after plating and collected 24 hours after the second transfection. For inducible TCF4 experiments, cells were treated with 2 ng/mL Doxycycline (Sigma-Aldrich, Cat# D9891) every 48 hours and collected 96 hours after plating. For ARV-771 experiments, cells were treated with 50 nM ARV-771 (Bioconnect, Cat#HY-100972) 24 hours after plating and collected 96 hours after plating. RNA was extracted using the RNA NucleoSpin extraction kit (Macherey&Nagel) according to the manufacturer’s instructions.

The RNA integrity was monitored using Bioanalyzer analysis. 5 ng of RNA per sample was reverse-transcribed and amplified using a modified version of the SMARTseq2 protocol, previously described in Rambow *et al.*^38^. Prior to generating sequencing libraries using the NexteraXT kit (Illumina, Cat#FC-131-10), cDNA profiles were monitored using the Bioanalyzer. Sequencing was performed on a Illumina Nextseq500 platform. Differential gene expression analyses were executed using the DeSeq2 pipeline.

### Geneset enrichment analysis

Geneset enrichment analysis was performed using GSEA 4.1.0. Briefly, approximately 3000 DEGs (si *TCF4* vs si Ctrl) were ranked by log2FC and the overlap with the following gene sets was estimated (MsigDB ID: M5930, M983 and M518).

### Western blotting

Harvested cell culture pellets were resuspended in protein lysis buffer (25 mM HEPES pH 7,5; 0,3 M NaCl; 1,5 mM MgCl2; 2 mM EDTA; 2 mM EGTA; 1 mM DTT; 1% Triton 0; 10% glycerol; phosphatase/protease inhibitor cocktail), incubated on ice (10min) and centrifuged at 14000 rcf for 15 minutes at 4°C. Equal amounts of protein, quantified using aLife Technologies Qubit 2.0 instrument were run on 4-12% Bis-Tris Plus Bolt gels (ThermoFisher Scientific) and transferred to a nitrocellulose membrane with an iBlot dryblot system (ThermoFisher Scientific). Membrane blocking (5% milk/TBS-0,2%Tween) was followed by incubation with the appropriate primary antibodies and HRP-conjugated secondary antibody. Signals were detected by enhanced chemiluminescence on Amersham hyperfilm. Antibodies that were used are the following:

|  |  |  |  |
| --- | --- | --- | --- |
| Rabbit polyclonal anti-MITF | Sigma-Aldrich | Cat# | HPA003259; RRID: |
| AB_1079381 |  |  |  |
| Rabbit monoclonal anti-TCF4 | Abcam | Cat# | ab217668; RRID: |
| AB_2714172 |  |  |  |
| Rabbit monoclonal anti-GAPDH | Cell Signaling | Cat# | 2118; RRID: |
| AB_561053 |  |  |  |
| Goat polyclonal anti-Rabbit IgG-HRP | Cell Signaling | Cat# | 7074; RRID: |
| AB_2099233 |  |  |  |

### Colony formation assay

Cells were grown to near confluency on 12-well plates and treated on the following day with 2 μg/mL Doxycycline (Sigma-Aldrich, cat# D9891) or vehicle. 48 hours after plating, cells were treated with the indicated drug combinations for four days. Cells were washed once with DPBS, stained with crystal violet (1% crystal violet w/v, 35% methanol v/v) for 15 minutes, washed with DPBS again and rinsed with tap water. Three regions of interests were quantified using the ImageJ plugin ColonyArea^94^ to define the intensity percentage. Each sample was then normalized over the relative control well and statistical significance was assessed t-test (unpaired, two-tailed Student’s t-test).

### Co-culture experiments

HLA-matched peripheral blood mononuclear cells (PBMCs) and MM099 cells were grown in 5% CO2 at 37°C for two days in RPMI 1640 medium supplemented with 10% FBS, 3 μg/mL anti-CD3 antibody (ThermoFisher Scientific, Cat#16-0038-85), 5 μg/mL anti-CD28 antibody (ThermoFisher Scientific, Cat#16-0289-85), 100 ng IL-2 (ThermoFisher Scientific, Cat#PHC0027).

TCF4 was silenced in MM099 cells as described above. Upon TCF4 knockdown, approximately 2000 MM099 cells per well were plated in a 96-well plate. Eight hours after plating, 10000 activated PBMCs per well were added, alongside aforementioned activating proteins and CellEvent™ Caspase-3/7 Green Detection Reagent (1:5000, ThermoFisher Scientific, Cat#C10423). Cells were imaged using the IncuCyte ZOOM System (Essen Bioscience) and automated apoptosis measurements were obtained based in images taken at 2-hour intervals, for the duration of the experiment. Three biological replicates were averaged and normalized over the last time point of the control and statistical significance was assessed (paired, two-tailed Student’s t-test).

### OmniATAC-seq

Approximately 2*10^5^ MM057 cells were plated in a 6-well plate and treated with ARV-771 100 nM after 24 hours. 48 hours after plating, cells were collected. Nuclei of 50,000 cells were isolated and an Omni-assay for transposase-accessible chromatin using sequencing (OmniATAC-seq) was performed as described previously^95^. After final amplification, samples were cleaned up with MinElute (QIAGEN) and libraries were prepareded using the KAPA Library Quantification Kit (supplier). Samples were sequenced on an IlluminaNextSeq 500 High Output chip.

Briefly, reads were mapped to human genome (GRCh37) using STAR (2.7.1a-foss-2018a). Resulting BAM files were cleaned for duplicates using Picard (2.21.8-Java-1.8.0) and indexed. Mitochondrial reads were removed using SAMtools (1.9-20190828-foss-2018a) and BigWig files were created (deepTools/3.3.1-foss-2018a-Python-3.7.4). ATAC-seq peaks were identified and visualized using MACS2 peak calling (2.1.2.1-foss-2018a-Python-2.7.16) with single-end BAMPE parameters. Finally, BED files were created from broadPeak files using BEDTools (2.28.0-foss-2018a).

### RT-qPCR

MM057 cells were plated and treated with the specified dose of ARV-771, as described above 48 hours after plating, cells were collected, resuspended in RA1 lysis buffer using the RNA NucleoSpin extraction kit (Macherey&Nagel) and processed according to manufacturer’s instructions. RNA was quantified using a ThermoScientific NanoDrop 1000 and 500 to 2000 ng was reverse transcribed with a High-Capacity cDNA Reverse Transcription Kit (Life Technologies). qPCRs were run using the SensiFAST probe No-ROX kit (Bioline, Cat#BIO-86005) on a Roche Life Science LightCycler 384. Data processing with Biogazelle Qbase+ 3.1 software relied on normalization with a minimum of two reference genes. RT-qPCR primer sequences are the following:

hTCF4 Forward: ATGGCAAATAGAGGAAGCGG

hTCF4 Reverse: TGGAGAATAGATCGAAGCAAG

hACTB Forward: CTGGAACGGTGAAGGTGACA

hACTB Reverse: AAGGGACTTCCTGTAACAATGCA

hRPL13A Forward: CCTGGAGGAGAAGAGGAAAGAGA

hRPL13A Reverse: TTGAGGACCTCTGTGTATTTGTCAA

hSDHA Forward: TGGGAACAAGAGGGCATCTG

hSDHA Reverse: CCACCACTGCATCAAATTCATG

### Proliferation assay

Roughly 400 MM029 cells per well were plated in a 96-well plate and treated with ARV-771 300 nM and/or Dabrafenib 50 nM + Trametinib 10 nM after 24 hours. Cells were imaged using the IncuCyte ZOOM System (Essen Bioscience) and automated cell confluency measurements were made using images taken at 2 hour intervals, for the duration of the experiments. Five technical replicates were averaged and statistical significance was assessed (paired, two-tailed Student’s t-test).

### TCGA SKCM data analysis

The SKCM raw count matrix composed of 375 samples was downloaded from Firehose. Raw TCF4 counts were calculated per sample and samples were grouped based on their phenotype^36^. Furthermore, log2 transformed read counts of were compared in metastatic versus primary melanoma lesions. Correlation analysis between *MITF* and *TCF4* mRNA levels in TCGA_SKCM was performed using cbioportal^96, 97^.

### Matrigel invasion assay and quantification

The invasive capacity of melanoma cells was determined by Matrigel transwell invasion assays using 0.8 mm BD BioCoatMatrigel Invasion Chambers (Corning, Cat#354480), according to manufacturer’s guidelines. Briefly, *TCF4* expression was knocked down in MM099 cells as described above. Next, cells were starved overnight in FBS- and L-glutamine-deprived medium. Around 2*10^5^ cells were plated in each chamber (coated with 25 μg Matrigel) in FBS-deprived medium, while 10% FBS- and 2.5% L-glutamine-enriched medium was used in the wells placed in the lower chamber. Uncoated inserts were used as a control for proliferation. 24 hours after seeding, membranes were stained with crystal violet.

Non-invading cells remaining on the upper surface of the chamber were removed by scrubbing with a cotton-tipped swab. Three to four randomly selected images were acquired per well and the surface cells were counted with ImageJ. The surface occupied by invading cells was calculated relative to the total surface of the membrane. Experiments included biological triplicates and technical duplicates.

### CCLE data analysis

*TCF4* read counts were plotted for skin_melanoma cell lines from the CCLE cohort^98^ and correlated (Pearson correlation coefficient) with IC50s (μM) of BRAF-(PLX4720) and MEK-(AZD6244) inhibitors.

### Data availability

Raw sequencing reads of all scRNA-seq have been deposited in the European Genome-phenome Archive (EGA) under study no. EGAS00001006488. Requests for accessing raw sequencing reads will be reviewed by the UZ Leuven-VIB data access committee. Any data shared will be released via a Data Transfer Agreement that will include the necessary conditions to guarantee protection of personal data (according to European GDPR law). Processed data of the malignant treatment naïve subset and the spatial transcriptomic RNA-sequencing data are available upon request.

