## Supplemental Figures for "A TCF4/BRD4-dependent regulatory network confers cross-resistance to targeted and immune checkpoint therapy in melanoma"

Supplemental Figure S1

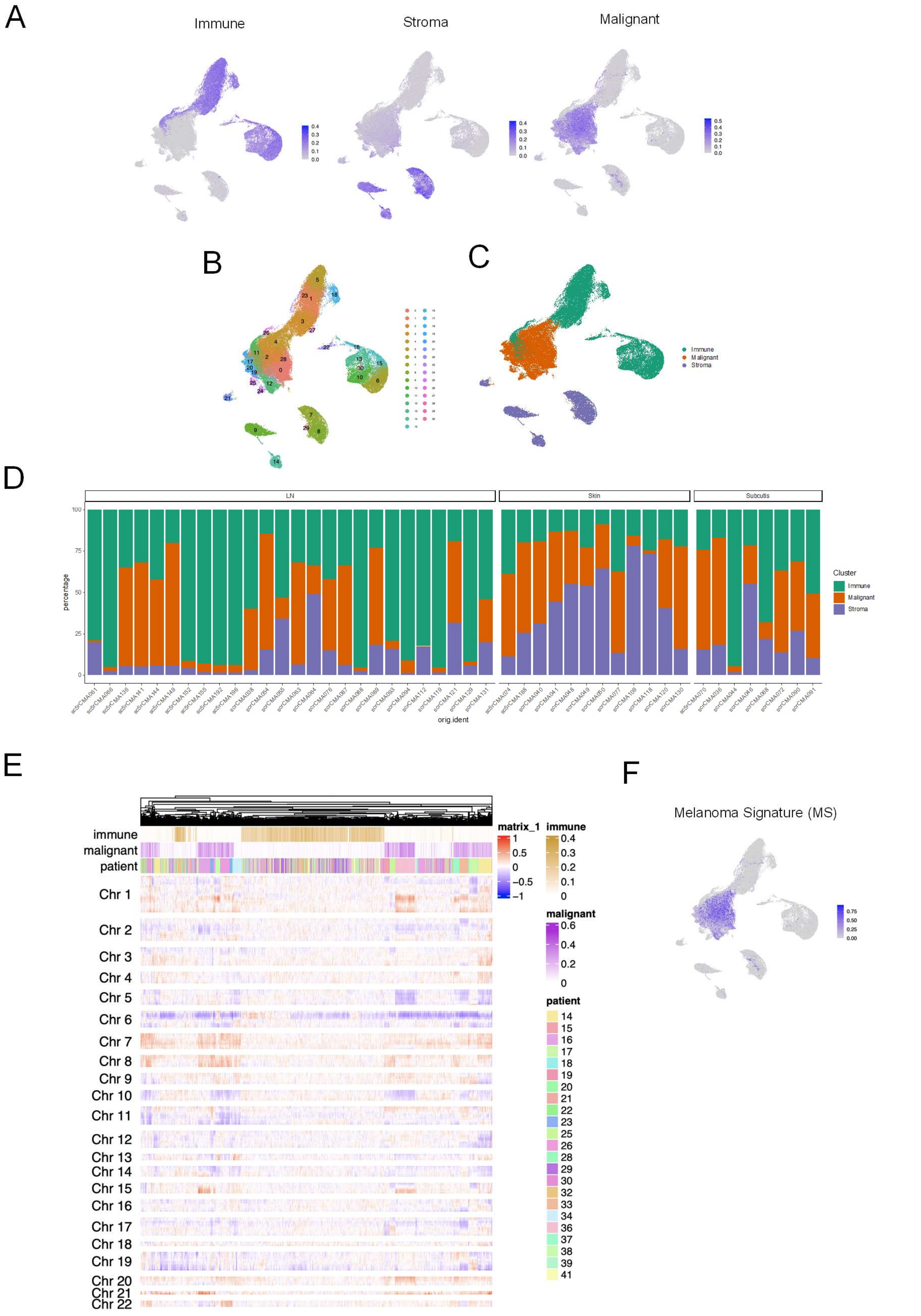

**Supplemental Figure S1: Cellular architecture of the entire melanoma ecosystem**

**A,** AUCell scores of the immune, stromal, and malignant signatures acquired from Jerby- Arnon *et al.*[22], projected on the global UMAP.

**B,** Unsupervised clusters on the entire tumour microenvironment.

**C,** Global UMAP plot of the entire tumour microenvironment at both time points.

**D,** Proportions of the three tumour microenvironment compartments across all the samples split by the biopsy site.

**E,** Heatmap of the inferred copy number variations across the patients. The top annotations also include the immune and malignant AUCell scores.

**F,** AUCell score of the Melanoma Signature (MS) projected on the global UMAP.

Supplemental Figure S2

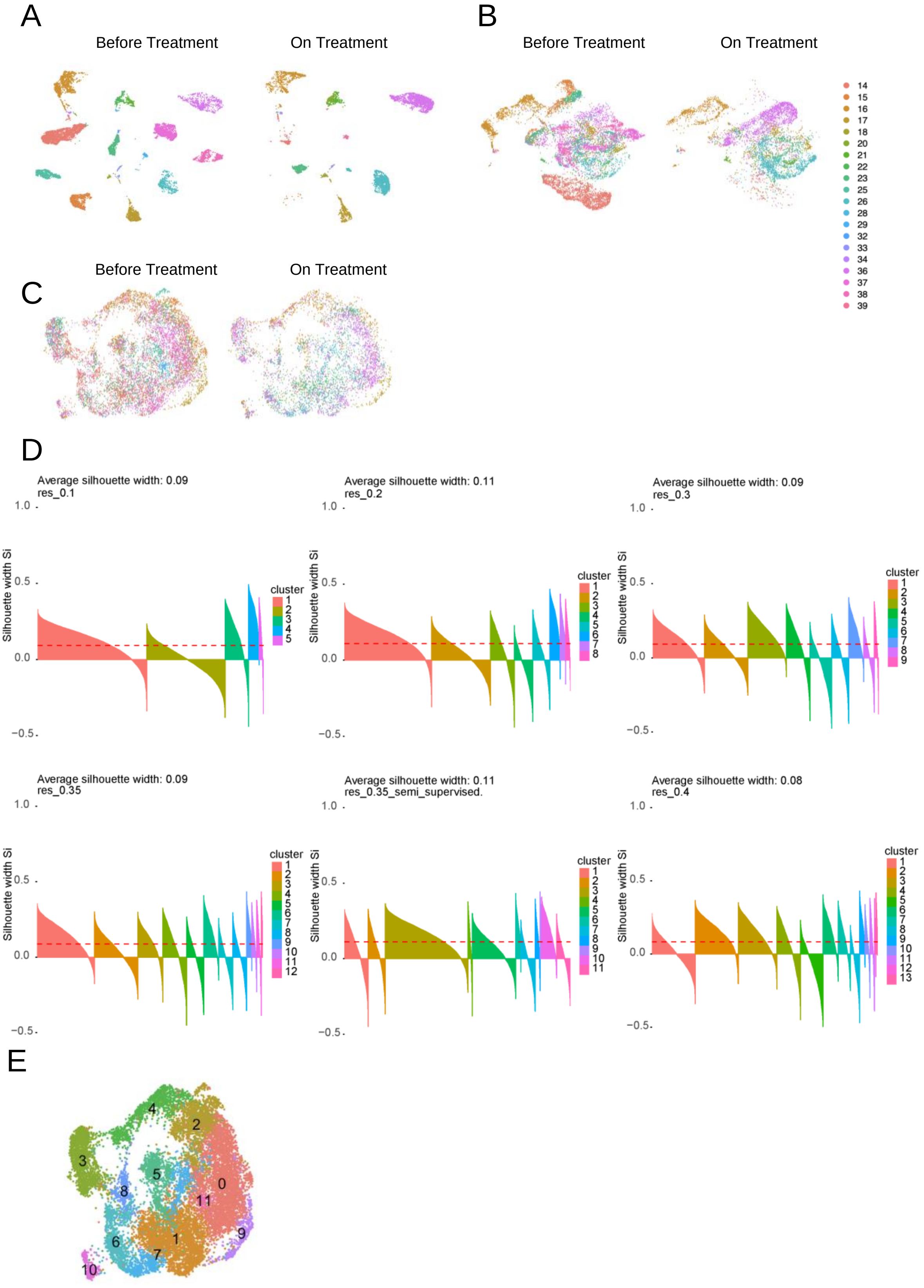

**Supplemental Figure S2: Clustering the malignant compartment**

**A,** Malignant cells clustered without sample ID regression.

**B,** Malignant cells clustered with sample ID regression.

**C,** Malignant cells clustered with data integration by sample ID using Harmony.

**D,** Silhouette scores for different clustering resolutions of the malignant cells, including semi-supervised clusters.

**E,** Unsupervised clustering of the malignant cells with resolution=0.35.

Supplemental Figure S3

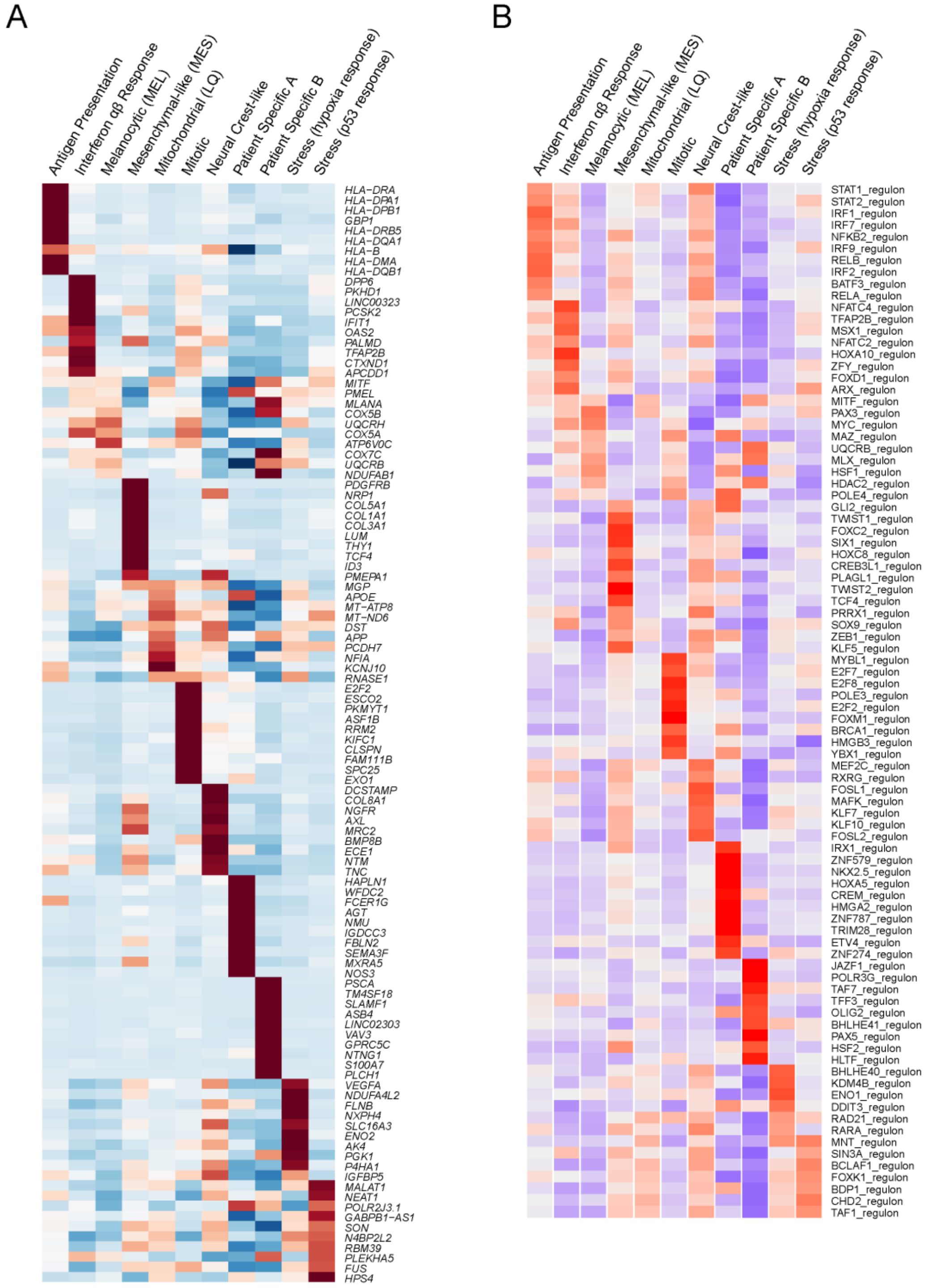


**Supplemental Figure S3: Differential gene expression and Regulon analysis of the melanoma cell states**

**A,** Heatmap of the selected top differentially expressed marker genes per malignant state.

**B,** Heatmap of the selected top differentially expressed regulons per malignant state.

Supplemental Figure S4

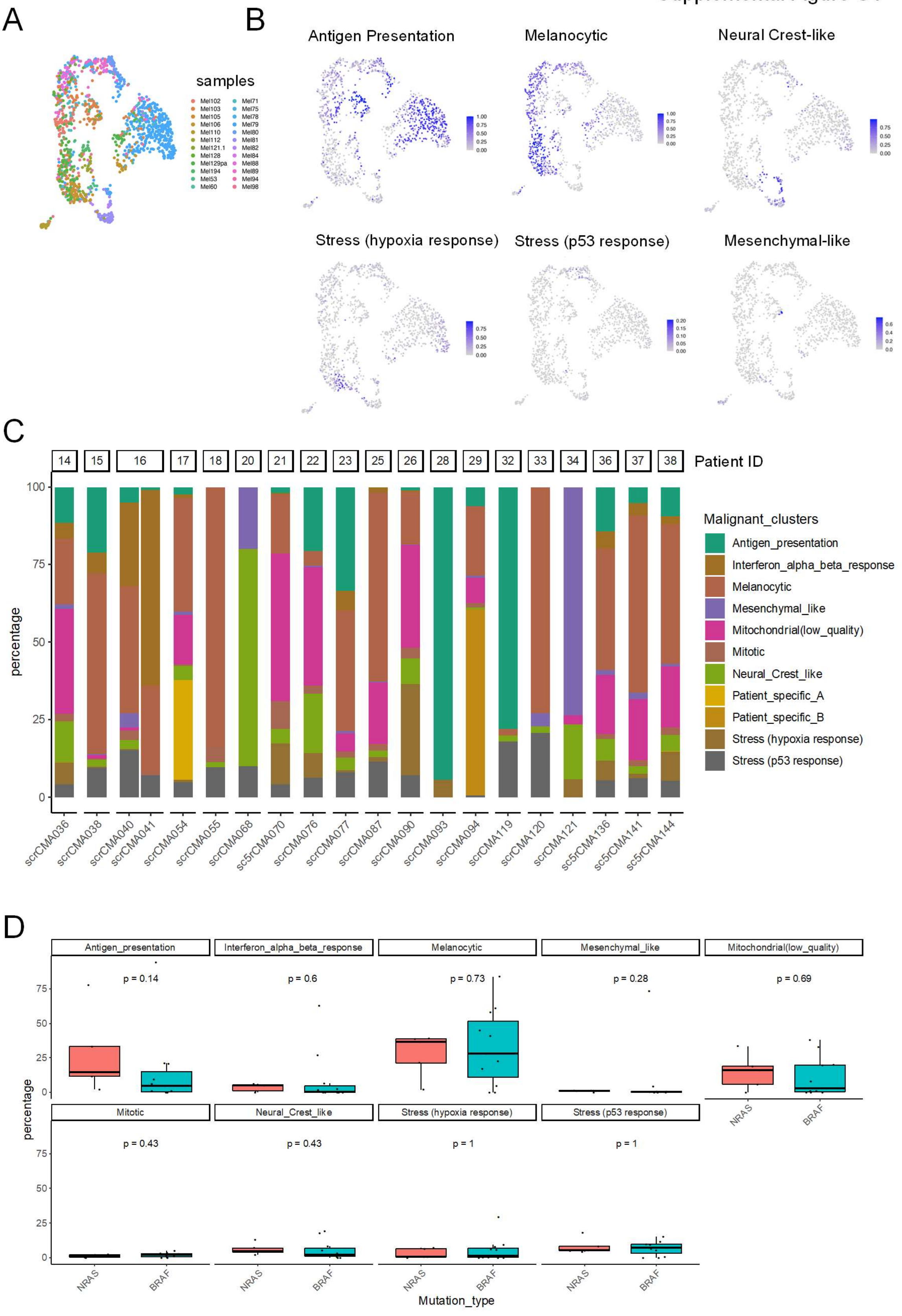

**Supplemental Figure S4: Monitoring melanoma cell states across various datasets**

**A,** UMAP of the malignant cells (integrated by sample ID with Harmony) from Jerby-Arnon *et al*.[22] study including both time points, coloured by sample ID.

**B,** Label transfer of the malignant states from our study onto malignant cells from Jerby-Arnon, *et al.*[22] using the standard Seurat pipeline.

**C,** Percentages of each of the malignant states per sample grouped by patient ID.

**D,** Percentages of each of the malignant states per molecular melanoma profile (*BRAF* and *NRAS mutant*) within the cohort described in this study (two-sided Wilcoxon test).

### Supplemental Figure S5

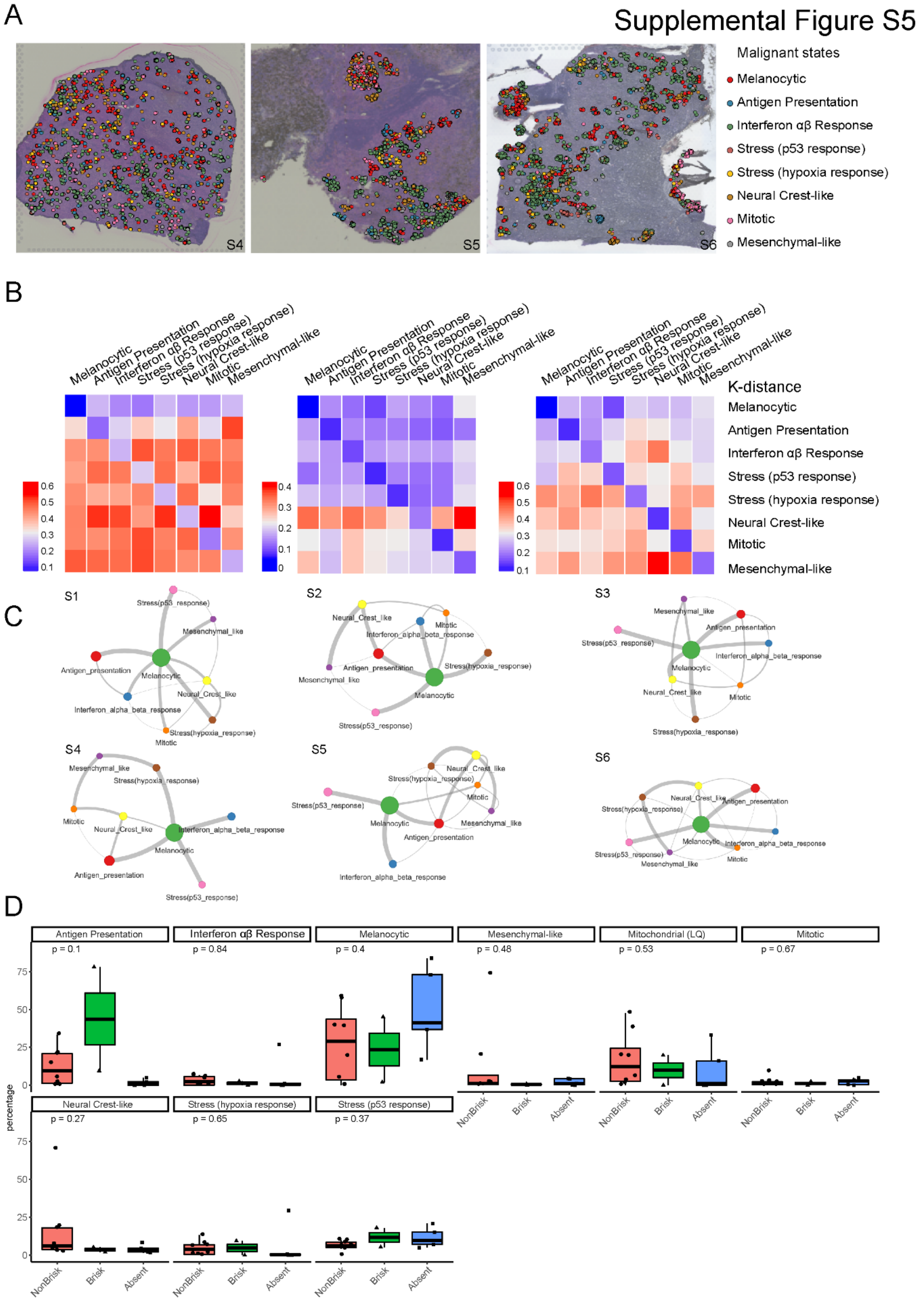

**Supplemental Figure S5: Mapping melanoma cell states in space**

**A,** Spatial transcriptomics of three additional treatment-naïve metastatic melanoma samples. Shown are malignant spots annotated per state.

**B,** Heatmaps of the k-distance calculated between the query state and every other cell state (rows). Note that as the k-distance metric is not normalized to the number of cells, comparisons can only be made within rows.

**C,** Simplified spatial cellular proximity calculated using the Delaunay Triangulation (DT), summarised in the graph abstractions per sample.

**D,** Percentages of cells from each of the malignant state tested between tumours categorized based on the TILs infiltration as assessed by our pathologist (Kruskal-Wallis test).

Supplemental Figure S6

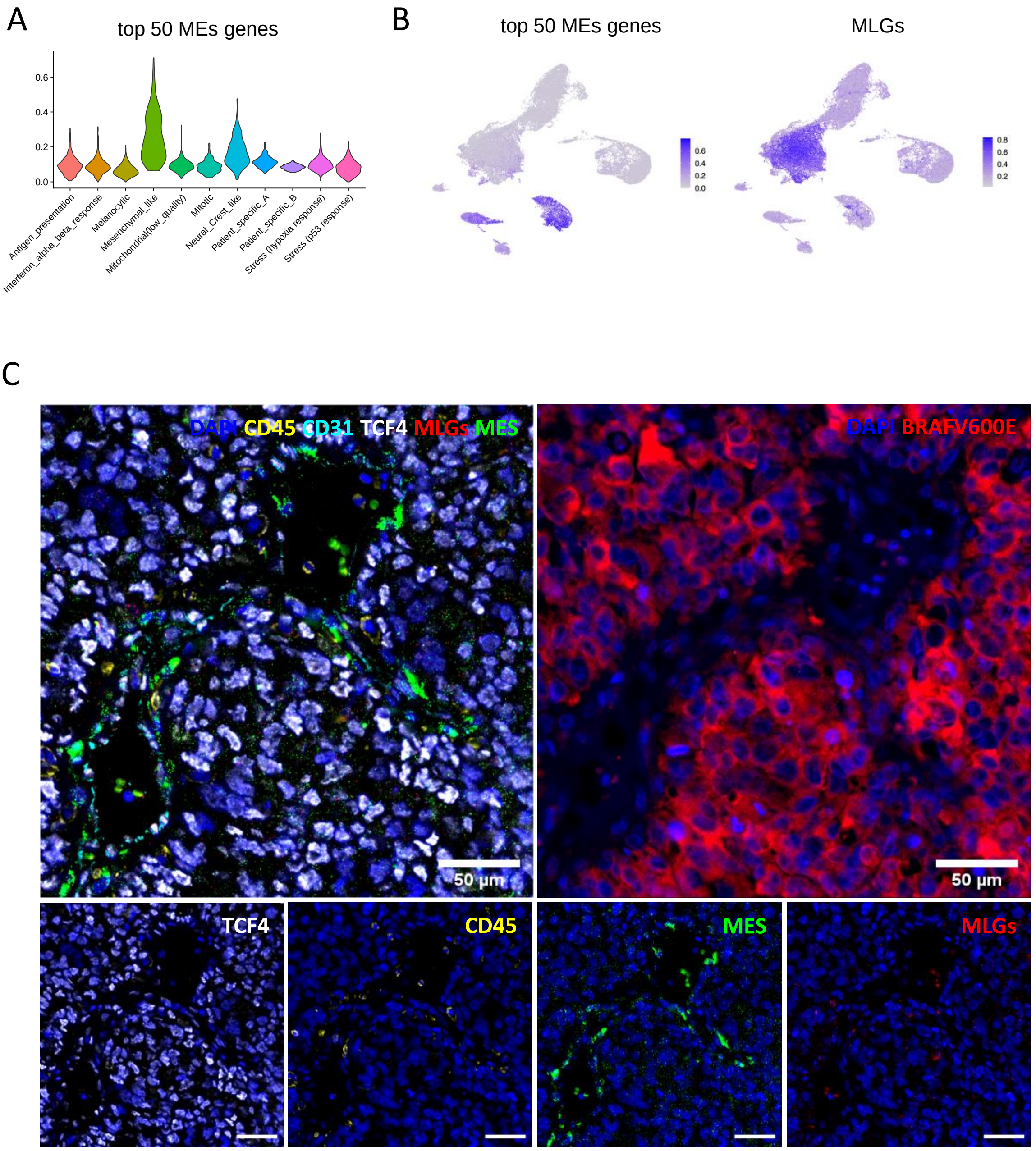

**Supplemental Figure S6: Developing methods to detect Melanoma MES cells**

**A,** AUCell score of the top 50 marker genes of the MES state plotted across malignant states.

**B,** AUCell score of the top 50 marker genes of the MES state and the MLGs plotted on the global UMAP plot (BT only).

**C,** Combined mIHC and mFISH image of a treatment naïve lymph node melanoma metastasis, harbouring a high proportion of MES cells. CD45, CD31 and TCF4 (white) protein stains are shown, whereas FISH of the four selected genes for the MLGs (*MITF*, *SOX10*, S100A1 and *CDH19; red)* and MES state (*DCN*, *TCF4*, *THY1* and *LUM* (green) are combined. A consecutive section was stained with a $BRAF^{V600E}$-specific antibody (right panel, red signal).

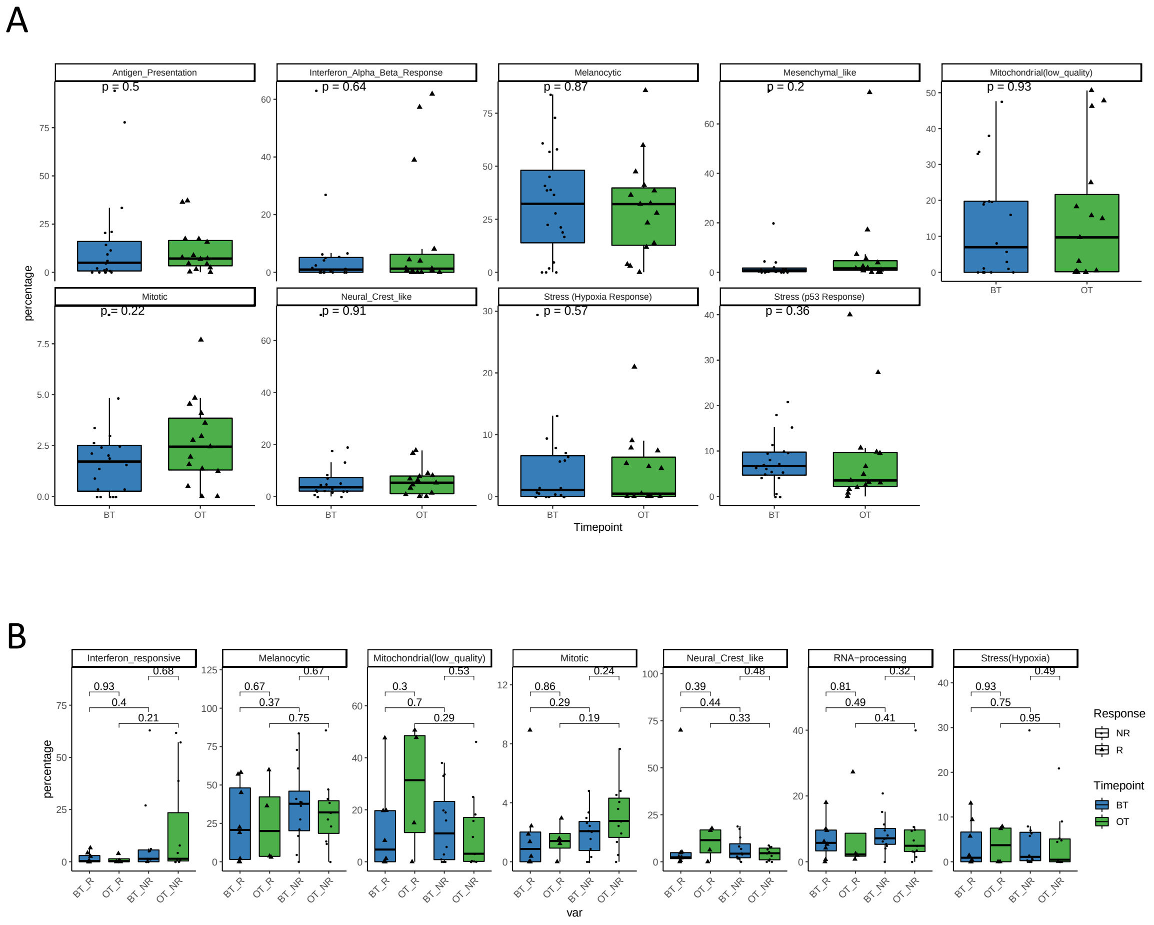


**Supplemental Figure S7: Melanoma cell states and response to ICB**

**A,** Percentage of malignant cells from each cell state out of all malignant cells per sample compared between the two time points (two-sided Wilcoxon test).

**B,** Percentage of malignant cells from each cell state out of all malignant cells per sample compared between R and NR at both time points (two-sided Wilcoxon test).

Supplemental Figure S8

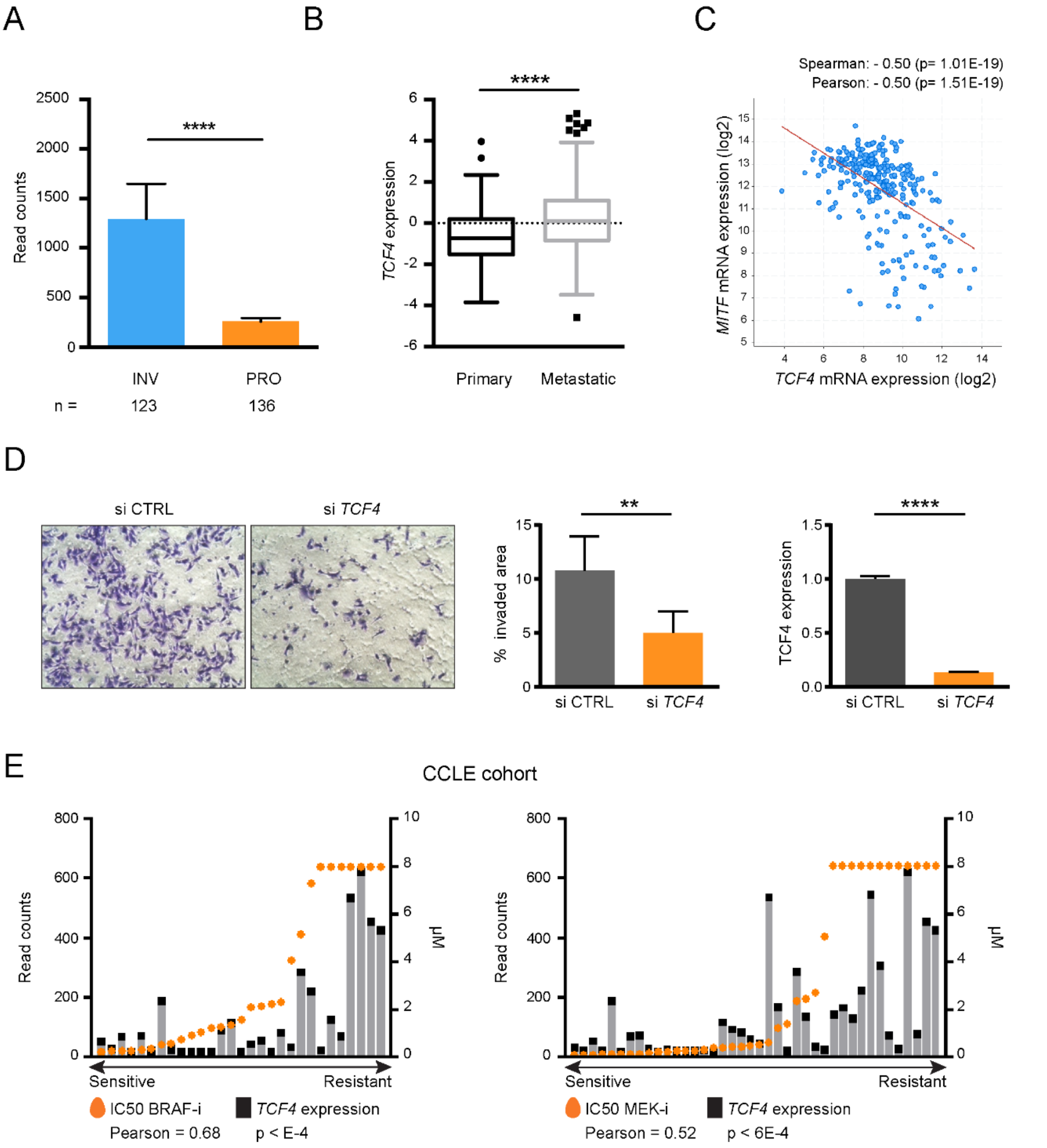

**Supplemental Figure S8: TCF4 contributes to the melanoma MES cell phenotypes**

**A,** *TCF4* expression (Read counts) in 375 TCGA SKCM samples (Mann-Whitney test, ****p<0.0001). INV, invasive; PRO=proliferative.

**B,** *TCF4* expression (log2 read counts) between Primary and Metastatic melanoma lesions (Mann-Whitney test, ****p<0.0001).

**C,** Correlation analysis between *MITF* and *TCF4* mRNA levels in TCGA_SKCM. Spearman and Pearson correlation are shown.

**D,** Left panel, matrigel-invasion assay upon silencing of *TCF4* in MM099 cells. Central panel, quantification (n=3 biological replicates; Mann-Whitney test, **p=0.0075). Right panel, relative *TCF4* expression upon silencing of *TCF4* (n=3 technical replicates, t-test, ****p<0.0001).

**E,** Correlation between *TCF4* expression and sensitivity (IC50, µM) to BRAF-inhibitor (PLX4720) and MEK-inhibitor (AZD6244) in the CCLE skin melanoma cell lines cohort. Pearson correlation is shown.

Supplemental Figure S9

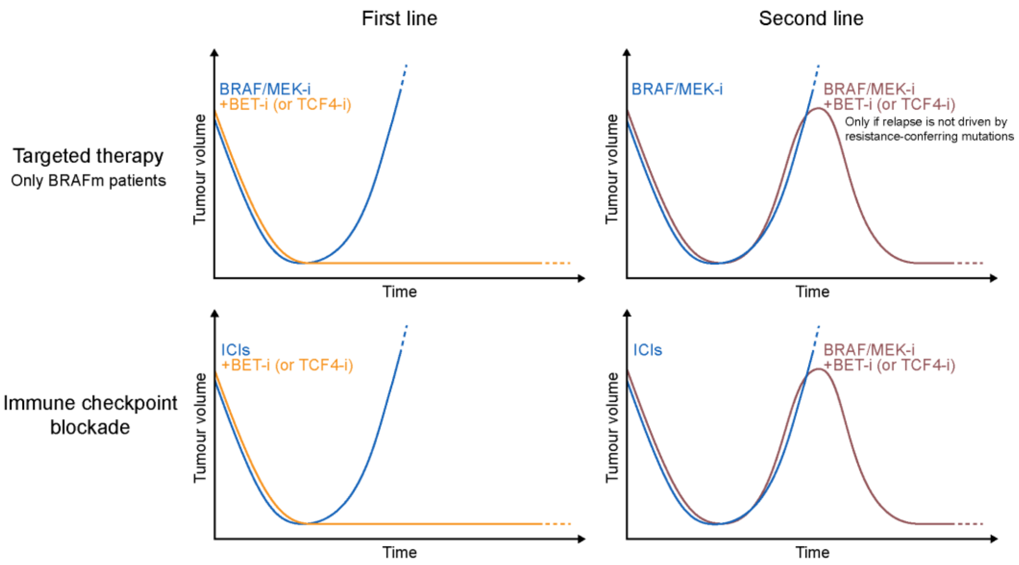


**Supplemental Figure S9: Schematic representation of clinical contexts in which BETi, or TCF4 targeting, may provide clinical benefit, through the targeting of melanoma MES cells.**
